# Biopsy-aware phylogenetic reconstruction from longitudinal single-cell copy number profiles

**DOI:** 10.64898/2026.10.14.759162

**Authors:** Jarosław Paszek, Agnieszka Mykowiecka, Krzysztof Gogolewski

## Abstract

**Motivation:** Longitudinal single-cell copy-number profiles provide information about cancer ancestry. Reconstructions that treat all observations as terminal leaves and disregard collection order cannot represent sampled ancestors or use their temporal relationships.

**Results:** We introduce reconstruction methods in the Cancer Tree Biopsy Framework (CTBF) that distinguish a copy-number profile from its separate observations across biopsies. Biopsy-guided reconstruction connects successive samples, defers ambiguous parent assignments, and completes the unresolved ancestry. Partial trees leave unresolved ancestral profiles unspecified. Closed-state trees assign every ancestor an observed profile. For comparison, pooled methods combine the profiles from all biopsies without using collection order. Across 100 independently simulated histories, the biopsy-guided partial methods attained higher mean rooted generalized Robinson–Foulds similarity than pooled neighbor joining and topology-only MEDICC2, using both complete and observed-state-restricted true trees. Under a fixed computational budget, they also attained higher mean similarity than NestedBD-Long at the two evaluated evolutionary depths. Convergence of its Markov chain Monte Carlo runs was not established within that budget. Classical neighbor joining and anticentral top completion gave closely similar partial-tree scores. Closed-state performance depended on evolutionary depth and the evaluation endpoint. An application to three longitudinal KTN152 biopsies demonstrated genome-scale feasibility on patient-derived integer copy-number estimates.

## 1 Introduction

Copy number alterations (CNAs), including focal and chromosome-scale gains and losses and whole-genome doubling, are pervasive in cancer and alter the dosage of oncogenes and tumor-suppressor genes [10, 29]. Single-cell DNA sequencing can resolve this variation as an integer copy-number profile (CNP) for each sampled cell, making it possible to study CNA evolution below the resolution of bulk clonal mixtures [16]. Phylogenetic reconstruction from CNPs remains challenging because the same locus may be altered repeatedly, and distinct event histories may reach the same copy-number state [4, 14]. Consequently, the same CNP state can correspond to distinct evolutionary occurrences.

Longitudinal biopsies provide information that is absent from a cross-sectional collection. Each biopsy is an incomplete observation of the tumor at one collection time, yet its order constrains ancestry between sampled occurrences. A CNP may also be observed at several times.

Collapsing these observations into one taxon would discard the distinction between recurrent or persistent occurrences, whereas forcing every observation to be a terminal leaf would preclude sampled ancestral relationships. Reconstruction from longitudinal CNPs therefore requires both CNP-state identity and biopsy-indexed occurrence identity.

Temporal information has previously been incorporated under several data models. CALDER constrains clone-tree inference from longitudinal bulk variant-allele frequencies [20]. LACE 2.0 infers a clonal tree from single-cell mutation profiles and constructs a longitudinal representation by duplicating clones across sampling times [1]. Phyllochron uses a supplied perfect-phylogeny clone tree, single-cell read counts, and sampling times to infer longitudinal cell-to-clone assignments by maximum likelihood [11]. Outside cancer phylogenetics, SeqTrack restricts direct ancestry among dated sampled genotypes and infers a minimum-cost branching [12]. NestedBD-Long is the closest recent method by input type. It uses binned single-cell CNPs and actual collection times to infer a Bayesian birth–death phylogeny with time-scaled branches and mutation rates [17]. We study a combinatorial setting in which biopsy order constrains ancestry, observed states may label internal nodes, and recurrent states retain distinct time-indexed occurrences.

Copy-number phylogenetic methods address complementary aspects of the problem. MEDICC jointly addresses allelic phasing, event-based dissimilarity, phylogenetic reconstruction, and ancestral-state inference [24]. MEDICC2 extends minimum-event reconstruction to include whole-genome doubling and uses neighbor joining for topology inference [14]. The `cnp2cnp` study introduces an event model allowing multi-copy gains and losses and evaluates its dissimilarity with classical neighbor joining [4]. Lazac reconstructs CNP phylogenies by parsimony under a zero-agnostic copy-number transformation model [22]. Sparse Rooted Neighbor Joining constructs rooted trees from a sparse subset of pairwise dissimilarities [30]. These approaches address event modeling and tree construction from CNPs. Our comparisons additionally examine the use of ordered biopsies.

Tree formalisms provide complementary precedents for the output representation. The copy-number-tree model assigns a CNP to every vertex [9]. Semi-labeled trees permit taxon labels at internal vertices and unlabeled internal vertices, while multiset-labeled trees permit repeated labels [6, 15]. Generally labeled Family-Joining trees and MEDALT lineage trees further show that an observed taxon or cell can occupy a parental position [13, 25]. In its MEDALT benchmark, MEDICC2 replaces an internally positioned observed cell by a latent parent and reattaches the observation through a zero-length edge [14]. In LACE 2.0, temporal duplication can also introduce an unobserved earlier instance of a parental clone [1]. Our missing-parent operation introduces an earlier occurrence carrying the child’s CNP state when the preceding biopsy supplies no unique plausible parent. This copy-up step preserves the later observation and carries the unresolved assignment to the next earlier transition. We formalize two corresponding output classes. A CNP-tree labels every vertex, whereas a partial CNP-tree allows unresolved internal vertices to remain unlabeled. Both classes permit the same CNP state to label distinct evolutionary occurrences.

Existing single-cell CNA simulators address broader biological and technological aims. CNAsim and SimSCSnTree can generate sequencing data, whereas CINner and SISTEM model population dynamics and selection [27, 19, 7, 28]. To evaluate reconstruction under controlled sampling, we use a minimal task-matched simulator that provides exact histories of genotype-state occurrences and ordered biopsy samples.

### Contributions

We introduce the Cancer Tree Biopsy Framework (CTBF) for reconstructing CNA evolution from longitudinal single-cell CNP observations. Our contributions are fourfold.

1. We formulate longitudinal CNP-tree reconstruction by separating CNP states from biopsy-indexed occurrences and by defining partial and closed-state output classes with recurrent state labels.
2. We develop a biopsy-guided connect–then–complete method that fixes supported relationships between consecutive biopsies, defers ambiguous assignments through a distinct copy-up rule, and then completes the unresolved top as either a partial or closed-state tree. Supplementary Methods 9.10 additionally define matched global temporal and no-time constructions.
3. We evaluate both output classes using a task-matched simulator and compare the proposed methods with pooled `cnp2cnp`–NJ, closed-state agglomerative baselines, topology-only MEDICC2, and finite-budget NestedBD-Long. The partial-tree comparison evaluates both complete and observed-state-restricted true trees and examines the contribution of top completion.
4. In an independently generated, prospectively specified simulation study, biopsy-guided partial methods attain higher mean similarity under both true-tree targets, with closely similar scores for classical NJ and anticentral top completion. When all ancestors must reuse observed profiles, performance depends on evolutionary depth and the evaluation endpoint. An application to three longitudinal biopsies from KTN152 demonstrates genome-scale feasibility on patient-derived CNP estimates.

## 2 Problem formulation

A tree *T* is a finite, connected, acyclic graph with node set *V*_*T*_ and edge set *E*_*T*_. A *rooted* tree is a pair (*T*, `root`(*T*)) where `root`(*T*) ∈ *V*_*T*_ is a distinguished node called the *root*. Viewing *T* as a directed acyclic graph obtained by orienting every edge away from `root`(*T*), the root is the unique node of indegree 0 and every other node has indegree 1. For *a, b* ∈ *V*_*T*_ we write *a* ⪯ *b* if *b* lies on the (unique) shortest path from `root`(*T*) to *a*. If *a* ⪯ *b* and *a ≠ b*, then *a* is a *descendant* of *b* and *b* is an *ancestor* of *a*, and we write *a* ≺ *b*. When the underlying tree must be made explicit, we write ⪯_*T*_ and ≺_*T*_ for these relations. A node with no descendants is a *leaf*, and all other nodes are *internal*. We denote the set of leaves by *L*_*T*_ ⊆ *V*_*T*_. The depth of a node *v* ∈ *V*_*T*_ is the number of edges on the unique path from `root`(*T*) to *v*. The height *h*(*T*) is the maximum node depth. For an integer *i* ≥ 0, *T* |_≤*i*_ is the rooted subtree induced by nodes of depth at most *i*.

### 2.1 Evolutionary histories and CNP-trees

Fix a common segmentation into an integer number *m* ≥ 1 of genomic bins and let 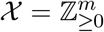 be the space of total copy-number profiles (CNPs) on that segmentation. For a finite, nonempty set of distinct profiles *P* ⊆ *X*, choose a set Σ_*P*_ of |*P*| state labels and fix a bijection ι_P_ : *P* → Σ_*P*_. Hence each state label *s* ∈ Σ_*P*_ identifies the exact profile 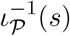. For a nonempty subset *P*^′^ ⊆ *P*, we retain its inherited labels ι_P_(*P*^′^) and use the restricted bijection 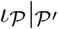.

#### Definition 1

(Evolutionary history tree). An *evolutionary history tree over P* is a pair (*T, p*_*T*_), where *T* = (*V*_*T*_, *E*_*T*_) is a rooted tree and *p*_*T*_ : *V*_*T*_ → *P* is a surjective profile assignment. Each node *v* ∈ *V*_*T*_ represents an evolutionary occurrence of the CNP *p*_*T*_ (*v*), and each directed edge (*u, v*) ∈ *E*_*T*_ represents a direct evolutionary transition from the occurrence represented by *u* to that represented by *v*.

The bijection ι_*P*_ gives the equivalent state-labeling function

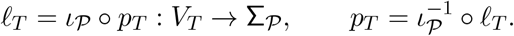

More generally, for a state-labeling function *ℓ* defined on a set *V*_ℓ_ ⊆ *V*_*T*_ with codomain Σ_*P*_, define 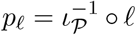. Throughout this paper, state-labeling functions need not be injective. Thus distinct nodes may represent distinct evolutionary occurrences of the same CNP. When the context is unambiguous, we write Σ, ι, and *ℓ* without subscripts.

#### Definition 2

(CNP-tree). A *copy-number–profile tree* (CNP-tree) over *P* is a pair (*T, ℓ*), where *T* = (*V*_*T*_, *E*_*T*_) is a rooted tree, *V*_ℓ_ = *V*_*T*_, and *ℓ* : *V*_ℓ_ → Σ_*P*_ is surjective.

When the CNPs of ancestral occurrences are unresolved, the corresponding nodes remain unlabeled. We represent such an evolutionary history by restricting the domain of the state-labeling function to the nodes whose CNPs are resolved.

#### Definition 3

(Partial CNP-tree). A *partial CNP-tree* over *P* is a pair (*T, ℓ*), where *T* = (*V*_*T*_, *E*_*T*_) is a rooted tree, *L*_*T*_ ⊆ *V*_ℓ_ ⊆ *V*_*T*_, and *ℓ* : *V*_ℓ_ → Σ_*P*_ is surjective.

A CNP-tree labels every node, whereas every unlabeled node of a partial CNP-tree is internal. We call such an unlabeled internal node a *latent node*. In both classes, surjectivity ensures that every profile in *P* is assigned to at least one node. These definitions are related to copy-number trees [9], semi-labeled trees [6], and multiset-labeled trees [15]. The distinctions among these formalisms are stated in Supplementary Methods 8.

### 2.2 Longitudinal biopsy samples

Let *T* ^★^ = (*T* ^★^, *ℓ*^★^) be a CNP-tree over *P* representing the true evolutionary history. We call *T* ^★^ the *true tree*. Let *q* ≥ 2 be an integer and let

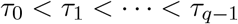

be the biopsy collection times. A *longitudinal biopsy sample* is an ordered family

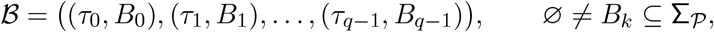

where *B*_*k*_ is the set of distinct CNP-state labels observed at time *τ*_*k*_. Because *B*_*k*_ is a set, multiplicities of a state within one biopsy are not part of the sample.

Define the observed state set and the set of time-indexed observations by

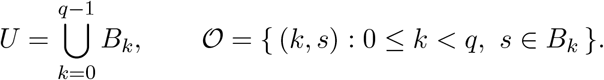

If the same state *s* is observed at distinct collection times, it occurs once in *U* but gives a distinct element (*k, s*) of *O* at each such time.

The sample *ℬ* is compatible with the true tree if there exists an injective sampling map

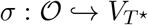

such that, for all (*k, s*), (*j, t*) ∈ *O*,

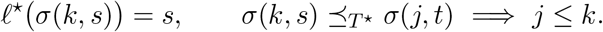

Thus time-indexed observations correspond to distinct occurrences, and ancestry among those occurrences is consistent with collection order. The true tree may also contain unobserved nodes, including nodes labeled by states in Σ_*P*_ *\U*.

### 2.3 CNP dissimilarities

The observed-profile map is *ξ* :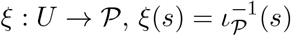. A *dissimilarity matrix* aligned with *U* is a matrix

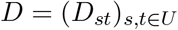

with finite entries satisfying *D*_*st*_ ≥ 0, *D*_*st*_ = *D*_*ts*_, *D*_*ss*_ = 0 for all *s, t* ∈ *U*.

We impose neither the identity of indiscernibles nor the triangle inequality. The matrix is indexed by states rather than time-indexed observations. Therefore all observations of the same state use the same row and column.

### 2.4 Reconstruction problems

The reconstruction input is *ℐ* = (*B, ξ, D*). Neither the true tree *T* ^★^ nor the sampling map *Σ* is part of *ℐ*.

A *reconstructed tree* for *ℐ* is a pair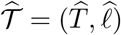, where 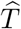 is a rooted tree and

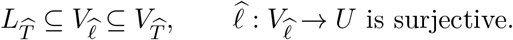

Its profile assignment on labeled nodes is 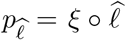. The reconstructed tree must admit an injective observation map

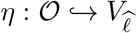

such that, for all (*k, s*),(*j, t*) ∈ *O*,

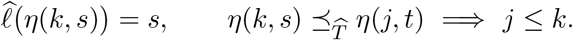

Thus distinct time-indexed observations are represented by distinct labeled nodes and respect collection order. The map *η* need not be unique when a state labels several nodes of 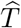.

#### Problem 1

(Partial CNP-tree reconstruction). *Given ℐ, compute a reconstructed tree* 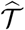. *Nodes in* 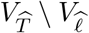 *represent unresolved internal occurrences*.

#### Problem 2

(Closed-state CNP-tree reconstruction). *Given ℐ, compute a reconstructed tree* 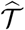 *satisfying* 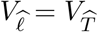. *Consequently, every node is assigned a state in U, and no ancestral CNP outside the observed profile set ξ*(*U*) *is introduced*.

The two problems define feasible output classes. Any criterion used to select among feasible trees is specified by the corresponding reconstruction method.

The input *ℐ* need not determine a unique true tree or the ancestor–descendant relations among observed occurrences, even when every ancestral state belongs to *U* (Supplementary Methods 8.3). A state first observed in a later biopsy may label an unobserved ancestor of an earlier sampled occurrence.

## 3 Reconstruction methods

We compare biopsy-guided reconstruction with pooled state-level and external copy-number comparators. Biopsy-guided methods construct feasible solutions to the partial and closed-state reconstruction problems. Pooled methods start from one occurrence per distinct state and may therefore collapse observations of a recurrent state. We classify every output by its node-labeling class. Exact randomization rules and auxiliary edge-weight conventions are specified in Supplementary Methods 9.

For states *s, t* ∈ *U*, define the irreversible-zero violation indicator

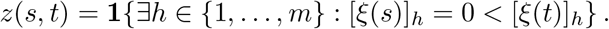

We call the direction *s* → *t zero-regain plausible* when *z*(*s, t*) = 0. This directional predicate is shared by the biopsy-guided, binary anticentral, and temporal constructions.

### 3.1 Reconstruction forests and top completion

#### Definition 4

(State-labeled rooted forest). A *state-labeled rooted forest over U* is a pair *ℱ* = (*F, ℓ*_*F*_), where *F* = (*V*_*F*_, *E*_*F*_) is a directed forest whose weakly connected components are rooted trees with parent-to-child edges, and *ℓ*_*F*_ : *V*_*F*_ → *U* is surjective. Its component roots are

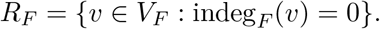

Nodes represent occurrences, so *ℓ*_*F*_ need not be injective.

#### Definition 5

(Top completion). Let *ℱ* = (*F, ℓ*_*F*_) be a state-labeled rooted forest. A *top completion* is a rooted tree 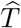 satisfying 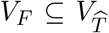 and 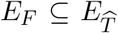. Let 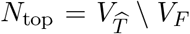. Both endpoints of every edge in 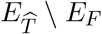 belong to *R*_*F*_ ∪ *N*_*top*_, and every top node is ancestral to at least one component root. Thus no edge of *F* is deleted, reversed, or subdivided. A *partial top completion* retains *ℓ*_*F*_ and leaves *N*_*top*_ unlabeled. A *closed-state top completion* extends *ℓ*_*F*_ to every node using only states in *U*. The resulting labeled tree is, respectively, a partial or closed-state CNP-tree. When one topology is used in both modes, the partial projection removes only the labels on *N*_*top*_. It changes neither the topology nor the labels of forest nodes.

A top completion can be expressed as an agglomerative procedure on the active component roots. A *reduction* merges two active components into one. At each reduction, a *pair-selection rule* chooses the two components. A topology-only reduction makes the selected roots children of an unlabeled ancestral node and therefore supports partial reconstruction. A closed-state reduction additionally applies a *parent-selection rule*, which selects an ancestral state or an existing ancestral root, and then merges the two components without assigning a state outside *U*. Pair selection and parent selection are distinct inference decisions.

### 3.2 Biopsy-guided bottom–top reconstruction

#### Definition 6

(Biopsy-guided bottom forest). A *biopsy-guided bottom construction* maps *ℐ* to a state-labeled rooted forest *ℱ*_*B*_ = (*F*_*B*_, *ℓ*_*B*_), a biopsy-index map *κ* :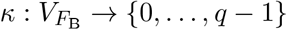, and an injective observation map *η* : 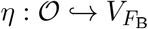 satisfying

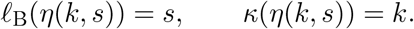

Every bottom edge (*u, v*) satisfies *κ*(*u*) *< κ*(*v*), every component root 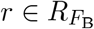 satisfies *κ*(*r*) = 0 and every additional occurrence is labeled by a state in *U*.

The bottom construction fixes inferred ancestry consistent with biopsy order and leaves a forest of lineages rooted at the earliest biopsy. Collection order does not order these roots relative to one another. A biopsy-guided method therefore applies a biopsy-order-independent top completion to *ℱ*_*B*_, using 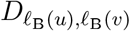 as the dissimilarity between component roots 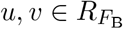. The choice between a partial and a closed-state output is made by the top completion. Because that completion adds ancestry only above roots at biopsy index 0, *η* remains a temporal-compatibility witness for the completed tree. Supplementary Proposition 4 proves this preservation property for the construction below.

Figure 1 summarizes the biopsy-guided bottom construction, top completion, and the binary pair–parent reduction shared by pooled and biopsy-guided reconstruction.

**Figure 1.**
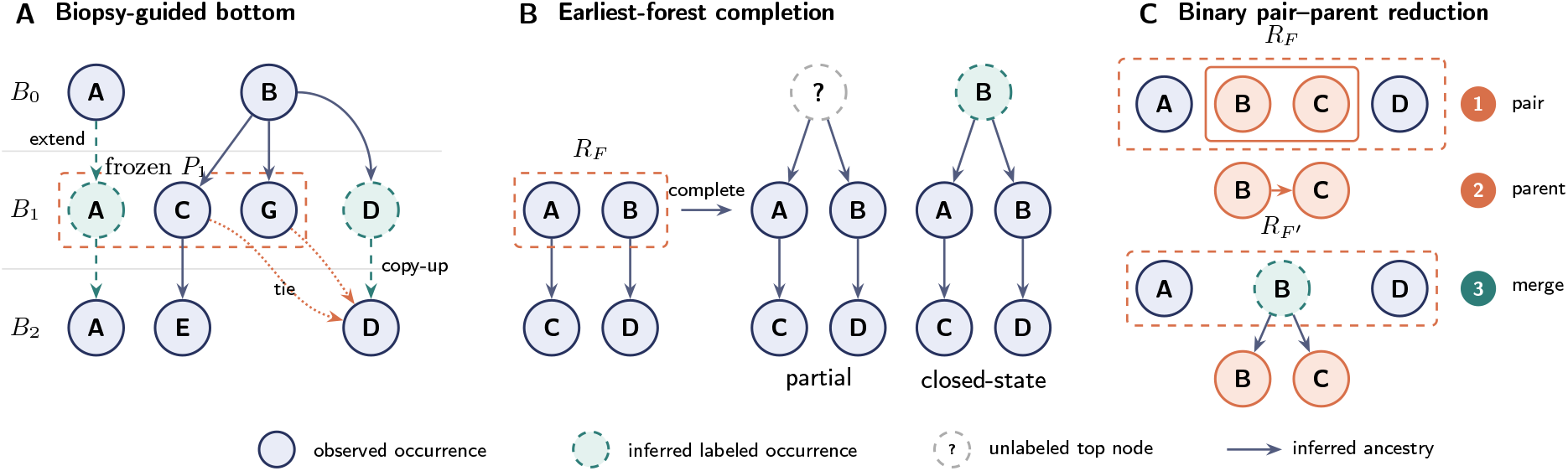
Biopsy-guided connect–then–complete reconstruction and shared binary pair–parent reduction. **(A)** Recurrent states are extended between their first and last observations before adjacent-biopsy assignment. The initialized candidate parents at biopsy index 1 form the set *P*_*1*_, which is frozen while the *B*_*1*_–to–*B*_*2*_ assignments are considered. Dotted arrows show tied candidate assignments, not retained ancestry edges. Deferred copies are added as a batch and participate as children in the next earlier transition. **(B)** A separate schematic two-component forest illustrates partial and closed-state completion. Both preserve all existing forest edges and labels. Only the partial completion leaves new top nodes unlabeled. **(C)** The upper rectangle contains the component roots *R*_*F*_ of the current forest *F*. Initially, the input is either the pooled singleton forest *ℱ*_*U*_, with one occurrence per observed state, or the biopsy-guided forest *ℱ*_*B*_. Existing descendant subtrees are omitted. Pair selection is followed by parent-state selection and a binary merge under a fresh occurrence of the selected state. The lower rectangle contains 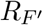, where *F* ^′^ is the forest after the merge. The middle arrow indicates the selected parent state. The binary anticentral rules instantiate this reduction for pooled reconstruction and biopsy-guided top completion. The partial projection removes only the labels of newly created top nodes.

#### 3.2.1 Adaptive/deferred bottom construction

The construction first extends every recurrent state across the interval between its first and last observation:

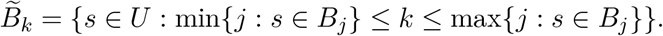

For each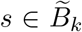, it creates an occurrence *v*_*k,s*_ labeled *s* at index *k*. The node *v*_*k,s*_ represents (*k, s*) when *s* ∈ *B*_*k*_ and is a level-extension occurrence otherwise. Initialize *F*_*B*_ with these vertices and no edges.

For *t* ∈ *{*1, …, *q* − 1*}* and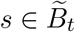, let 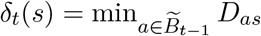 *D*_*as*_ and set

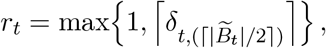

where *δ*_t,(j)_ is the *j*th order statistic of the indexed family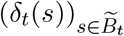, with repeated values retained. All radii are fixed before parent assignment. Transitions arepro cessed from *q* − 1 down to 1. At the start of transition *t* − 1 → *t*, define 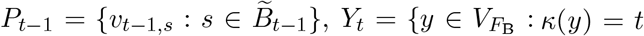, indeg 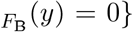. At that stage, *Y*_*t*_ contains both initialized occurrences at index *t* and copies propagated from the already processed transition *t* → *t* + 1. The set *P*_*t*−*1*_ is frozen throughout the transition. For each *y* ∈ *Y*_*t*_, the same-state occurrence in *P*_*t*−*1*_ is preferred. Otherwise, the parent is the unique minimum-distance *x* ∈ *P*_*t*−*1*_ satisfying 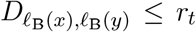 and *z*(ℓ_*B*_(*x*), *ℓ*_*B*_(*y*)) = 0. If no such unique minimizer exists, the decision is deferred. After all children have been considered, each deferred child *y* receives a fresh parent occurrence 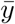 at index *t* − 1 with 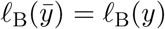. These copies are inserted as one batch, so none can parent another child in the same transition. Each instead participates as a child in the next earlier transition. Exact order-statistic, edge-weight, and bookkeeping conventions are specified in Supplementary Methods 9.3. The temporal condition applies to sampled occurrences rather than to state labels. A state observed only in a later biopsy may therefore label an unobserved occurrence on the path to an earlier sample. When no unique plausible parent is available, deferred copy-up creates such an earlier occurrence with the child’s state, allowing the construction to represent the state-*a* founder in Figure 4.

### 3.3 Pooled state-level input

Any top-completion rule can instead be applied to the *pooled singleton forest ℱ*_*U*_ = (*F*_*U*_, *ℓ*_*U*_), where

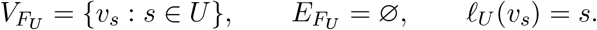

This biopsy-order-independent input contains one occurrence per state rather than per observation in *O*. Consequently, a recurrent state observed in several biopsies may be represented only once, precluding the injective observation map required by the reconstruction problems.

### 3.4 Binary anticentral top completion

Let *a*_*1*_, …, *a*_*k*_ be the active component roots, and let Δ_*ij*_ denote their current dissimilarities.

Each *c*_*i*_ ∈ [0, 1] is the current centrality score of *a*_*i*_. Initially, 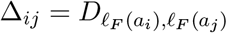, and *c*_*i*_ is the normalized inverse of the mean dissimilarity from *a*_*i*_ to the initial roots. Thus larger initial scores indicate smaller mean dissimilarities. At each reduction, the method selects the pair minimizing

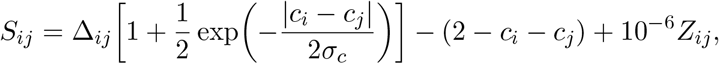

where *Σ*_*c*_ *>* 0 is the stabilized standard deviation of the current centrality scores, and the *Z*_*ij*_ are independent uniform random variables on [0, 1) drawn from the method’s seeded generator. The perturbation is applied to every active pair as part of its score. Selection favors small dissimilarities and low total centrality. The exponential factor imposes a larger distance penalty on pairs with more similar centralities.

For parent selection, define diploid deviation

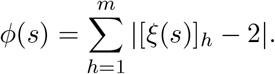

For selected roots *a*_*i*_, *a*_*j*_, the sole zero-regain-plausible direction is used when only one exists. If both directions are plausible, the state with smaller *ϕ* is ancestral. A tie in *ϕ*, or two implausible directions, is resolved in favor of the larger *c*, followed by a seeded exact-tie rule. The reduction creates a fresh parent occurrence carrying the selected parent state and makes both selected roots its children. The parent distance row is retained, the selected centralities are averaged, and reduction continues until one root remains. Exact initialization, stabilization, perturbation, merge, and update rules are given in Supplementary Methods 9.4.

Applied to *ℱ*_*U*_, this rule defines the pooled closed-state method *M*_*5*_. Applied to *ℱ*_*B*_, with the same initial root order and random seed, it defines two projections of one topology: the partial method *M*_*2*_ suppresses labels on *N*_*top*_, whereas the closed-state method *M*_*6*_ retains the copied parent-state labels. Their forest nodes, including level-extension and deferred-copy occurrences, are identical.

### 3.5 Comparators

Classical neighbor joining (NJ) selects pairs by the *Q* criterion and introduces unlabeled latent nodes [21]. Applied to *ℱ*_*U*_ with the `cnp2cnp` dissimilarity specified in Supplementary Methods 10.2, it defines the pooled partial baseline *M*_*0*_ [4]. Subdividing the final joining edge with an unlabeled synthetic root supplies the rooted representation used for evaluation. Root placement is a representational convention. Applying the same NJ completion to *ℱ*_*B*_ defines *M*_*4*_. It retains the adaptive/deferred bottom of *M*_*2*_ and replaces its anticentral top by classical NJ.

MEDICC2 provides an external comparator designed specifically for copy-number phylogenies [14]. It receives every member of *O* as an opaque taxon without its biopsy index and computes its own minimum-event dissimilarities and NJ topology. After removal of its artificial diploid outgroup, observed occurrences remain labeled leaves and internal nodes remain unlabeled. We therefore evaluate MEDICC2’s distance calculation and topology inference jointly as an external partial-tree pipeline.

NestedBD-Long uses observation CNPs and numerical collection times to infer a time-scaled birth–death phylogeny [17]. We compare its sampled-ancestor-aware topology with the partial reconstructions. The fixed-budget inference, tree summary and mapping of observation labels are specified in Supplementary Methods 11.7.

Two pooled closed-state comparators separate pair selection from parent selection. Method *M*_*1*_ selects the closest pair and copies the state of the member with smaller residual row sum. Method *M*_*3*_ selects pairs by the NJ *Q* criterion and directly orients the pair using fixed initial total-dissimilarity scores. Applying the latter completion to *ℱ*_*B*_ defines the biopsy-guided closed-state method *M*_*7*_. Exact recurrences, roots, and tie rules are given in Supplementary Methods 9.5–9.8. Computational bounds for all principal methods are given in Supplementary Methods 9.9. Supplementary Methods 9.10 additionally specify matched global temporal and no-time methods on *O*.

### 3.6 Method composition map

Table 1 identifies each method by its input representation, completion rule, and output-labeling class.

**Table 1.**
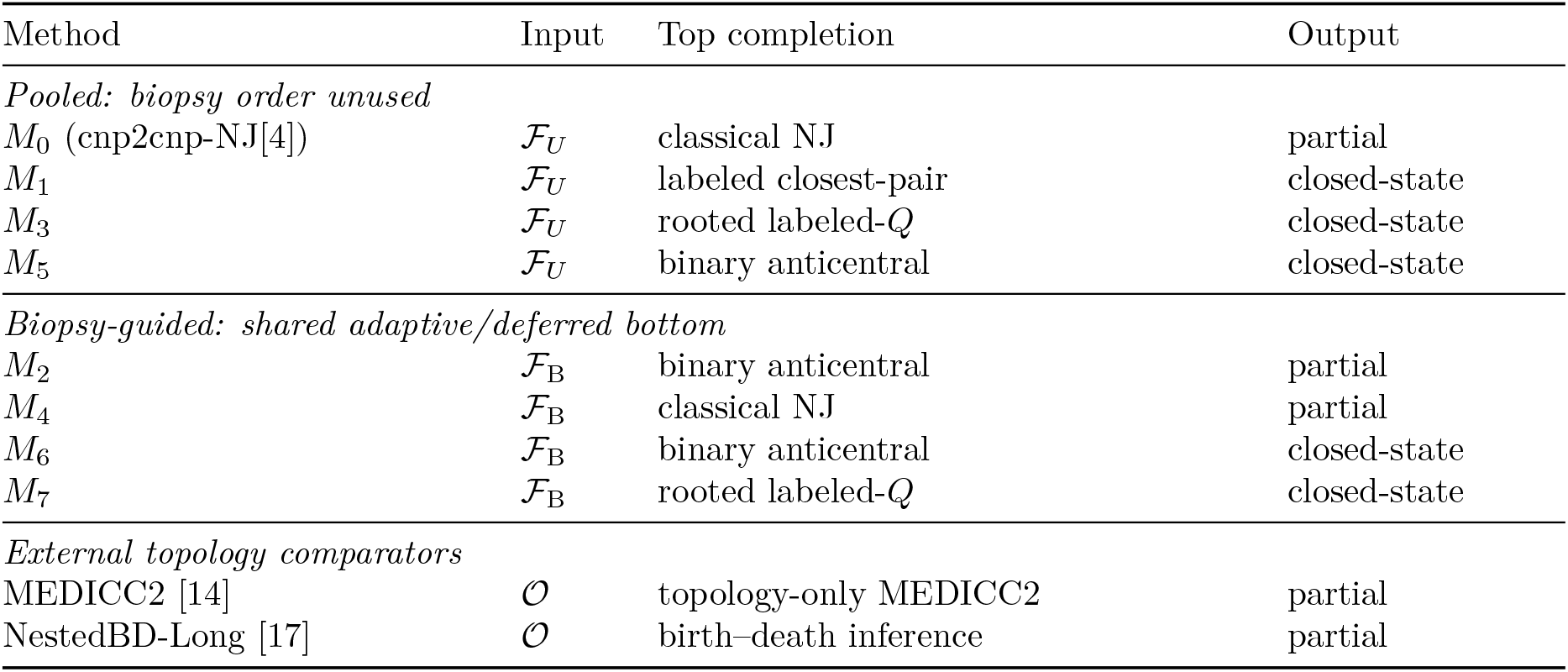
Composition of evaluated reconstruction methods and external comparators. NestedBD-Long additionally receives numerical collection times.

Methods *M*_*2*_ and *M*_*6*_ are partial and closed-state projections of one biopsy-guided reconstruction. Comparing *M*_*5*_ with *M*_*6*_ changes only the initial forest and its use of biopsy order while preserving the anticentral completion. Methods *M*_*3*_ and *M*_*7*_ analogously share rooted labeled-*Q* completion. Method *M*_*1*_ is the distinct raw-distance copied-parent baseline.

## 4 Experimental design

We evaluated the reconstruction methods in two complementary settings. Paired simulations provide known evolutionary histories for measuring recovery by CTBF, MEDICC2 and NestedBD-Long. A separate application to three longitudinal KTN152 biopsies assesses genome-scale execution on patient-derived CNP estimates without a truth-based accuracy endpoint.

### Synthetic design and inputs

The CTBF simulator generates discrete-generation true CNP-trees under chromosome-bounded singleton and interval gains and losses, retaining at most one occurrence of each viable CNP per generation. Generation size therefore counts distinct viable CNP states. These exact histories support controlled evaluation of reconstruction from sampled profiles. The Discussion addresses the biological and observational scope of this model. We independently generated 100 seeded histories to height 34. From each history, we formed three standard conditions by applying the same three-biopsy schedule to prefixes ending at heights 14, 24, and 34. We formed five additional conditions from the height-24 prefix. Three replaced the standard three biopsies with two, four, or five biopsies. Two retained three biopsies but used late or random sampling times. This yielded eight conditions per history.

For each condition, the minimum of both directed `cnp2cnp` costs between every pair of its observed states defined the symmetric input matrix *D* [4]. All CTBF methods received the same state-aligned matrix. MEDICC2 received the same occurrence profiles for the 300 standard conditions, but no biopsy indices, and computed its own dissimilarities. NestedBD-Long received the unmodified occurrence profiles and numerical sampling generations for the 200 height-14 and height-24 standard conditions. Each input was assigned two independently seeded runs of two million iterations. Its inference and topology-only evaluation are specified in Supplementary Methods 11.7. Table 3 gives the simulator parameters. Biopsy schedules, sampling, and dissimilarity construction are specified in Supplementary Methods 10.1–10.2. The sampling fraction specifies observation density among retained genotype-state occurrences.

### Evaluation

Before scoring, we require each true and inferred graph to be a nonempty rooted tree with parent-to-child edges. Every inferred state label must belong to *U*, and every state in *U* must occur at least once in the inferred tree. Invalid or incomplete outputs remain explicit failures. Ancestor–descendant F1 (AD-F1) compares unique ordered state-label ancestry relations and is the primary endpoint for closed-state reconstructions. It does not measure direct-edge recovery or correspondence between repeated occurrences. Its true relation set is restricted to states in *U*. Rooted generalized Robinson–Foulds (GRF) similarity compares multisets of subtree label multisets [18]. It preserves repeated labels and equal clusters and is the primary endpoint for partial reconstructions. It is also a complementary endpoint for closed-state reconstructions and compares the complete tree representations, so unobserved true states and tree-size differences contribute. Partial trees are scored in the rooted representations specified above, including the synthetic final-join root of *M*_*0*_, the tumor-subtree root retained for MEDICC2 and the sampled-ancestor-aware projection of NestedBD-Long. Exact metric definitions are given in Supplementary Methods 10.3. Larger values indicate better agreement for both scores.

A complementary evaluation target is the *trimmed true tree*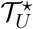. It retains every true occurrence of a state in *U* and the unlabeled branching nodes needed to connect these occurrences. GRF against this target uses the unchanged reconstructed tree (Supplementary Methods 10.3.3). We report full-truth and trimmed-truth GRF for the partial-tree comparison, with supplementary trimmed-truth analyses of biopsy number and closed-state reconstruction. The prespecified primary comparison retains the full true tree. AD-F1 uses the same observed-state ancestry relation set under both targets.

### Paired comparisons

Comparisons pair methods or sampling conditions within the same simulated history. A gain from additional biopsies is the score with more biopsies minus that with fewer, evaluated within each history. We report mean scores and paired differences with pointwise 95% bootstrap intervals. Resampling treats each independently simulated history, together with all its paired results, as one unit. For NestedBD-Long, the two successful run scores are averaged within each input before histories are resampled. The five-method partial-tree panel uses a common successful-history set at each height. Combined-height contrasts first average the height-specific differences within each complete history. Detailed tests, failure handling, and resource accounting are given in Supplementary Methods 10.4.

## 5 Synthetic reconstruction results

We evaluated reconstruction methods on 100 independently seeded histories that were not used for method development. All 800 prespecified input conditions, 6,400 internal reconstruction records, and 300 topology-only MEDICC2 records completed without failure. The NestedBD-Long extension attempted both height-14 and height-24 inputs from every history, with two runs per input. Of 400 runs, 397 succeeded. One posterior-consistency failure and two timeouts affected two height-24 inputs, leaving 100 paired inputs at height 14 and 98 at height 24. Scoring and paired inference follow Supplementary Methods 10.3 and 10.4. Square brackets in the text denote pointwise 95% bootstrap intervals obtained by resampling whole histories.

### Additional biopsies improve ancestry recovery

At height 24, mean closed-state AD-F1 increased between two and five biopsies for *M*_*3*_, *M*_*5*_, *M*_*6*_ and *M*_*7*_ (Figure 2A), while *M*_*1*_ declined. For *M*_*6*_, the mean increased from 0.73391 to 0.79928, giving a paired gain of 0.06537 [0.04978, 0.08198]. For *M*_*7*_, the corresponding gain was 0.08394 [0.06326, 0.10492]. These scores assess ancestry among all states observed in each condition.

**Figure 2.**
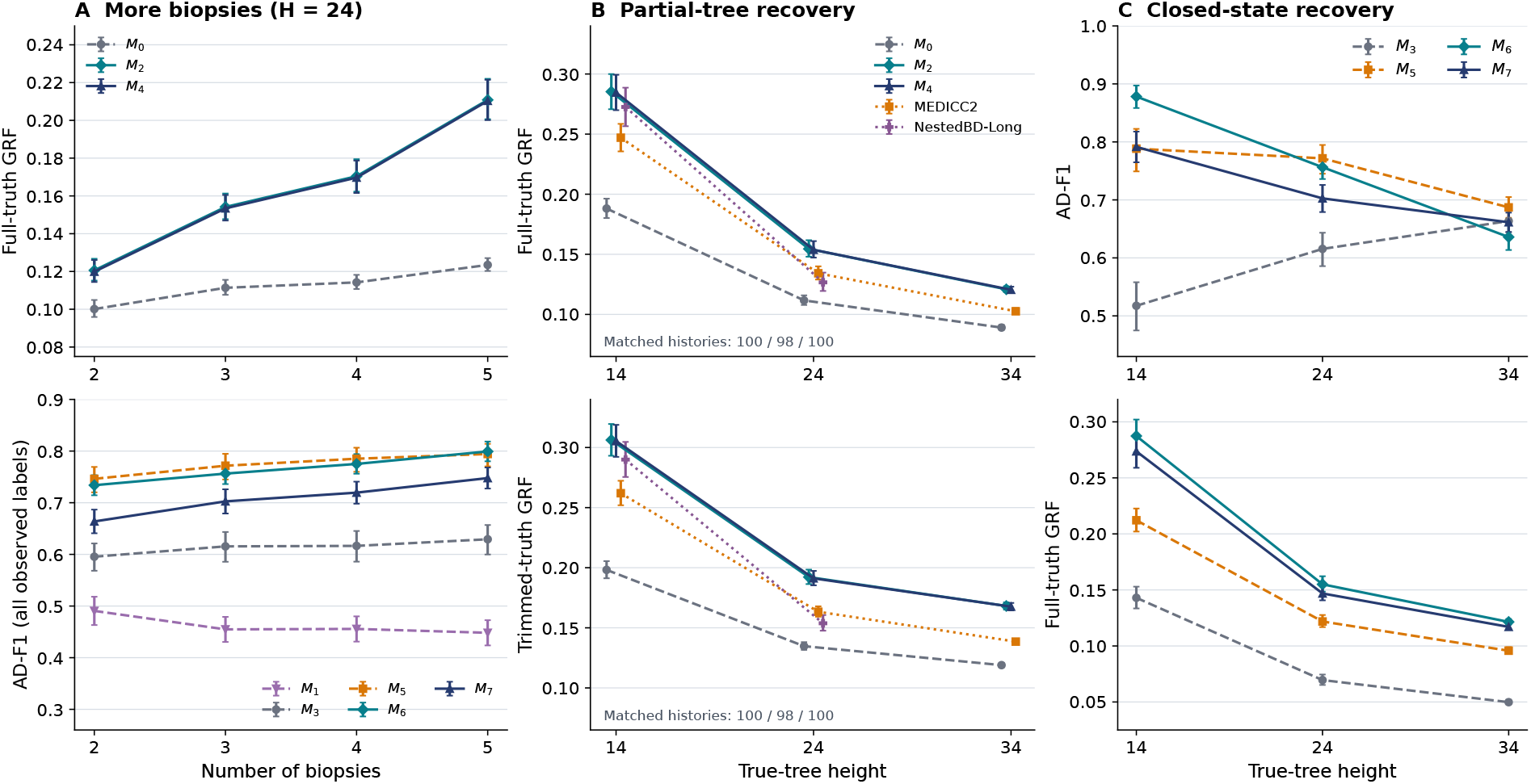
Reconstruction accuracy by biopsy number and evolutionary depth. **(A)** Mean scores for eight CTBF methods at height 24 with two to five biopsies. The upper plot shows full-truth GRF for partial trees from *M*_*0*_, *M*_*2*_ and *M*_*4*_. The lower plot shows ordinary closed-state AD-F1 using all labels observed in the corresponding condition. Its scoring label set grows with biopsy number. Solid, dashed and dotted curves denote biopsy-guided CTBF, pooled CTBF and external methods, respectively. **(B)** Partial-tree GRF against the full true tree (upper) and the trimmed true tree (lower). All available methods use the same 100, 98 and 100 successful histories at heights 14, 24 and 34, respectively. NestedBD-Long contributes the mean of two two-million-iteration run scores per input and has no height-34 point. Small horizontal offsets separate overlapping markers. **(C)** Mean AD-F1 (upper) and full-truth GRF (lower) for four closed-state methods. Panels A and C use all 100 histories. Error bars are pointwise 95% percentile intervals from 100,000 bootstrap resamples of whole histories. Partial-tree GRF retains occurrence multiplicities. External methods were assessed only under the standard sampling schedule. Supplementary Figure <u>7</u> gives the fixed-label AD-F1 control and trimmed-truth counterparts of panels A and C.

A complementary control scores only the states already observed with two biopsies, while retaining the additional observations as reconstruction input (Supplementary Figure 7A). Its mean gains were 0.09254 for *M*_*6*_ [0.07940, 0.10655] and 0.10206 for *M*_*7*_. Both exceeded the gains of their pooled counterparts with the same completion rule. Pooled *M*_*5*_ also gained 0.04714. Thus additional observations improved recovery even when the scored ancestry target was held fixed.

For partial trees, the mean full-truth GRF gains were 0.09023 for *M*_*2*_, 0.09044 for *M*_*4*_ and 0.02337 for *M*_*0*_. Against trimmed truth, the corresponding gains were 0.07796, 0.07868 and −0.00275. The biopsy-guided gains therefore persisted under both targets, whereas the pooled-NJ contrast changed sign. The trimmed target changes with the observed-state set, so these sampling contrasts do not hold the target fixed. Tables 7–9 and 14 give the descriptive biopsy-number comparisons.

### Biopsy guidance improves partial reconstruction

Methods *M*_*2*_ and *M*_*4*_ attained higher mean partial-tree similarity than pooled NJ and topology-only MEDICC2 at all three heights, and than NestedBD-Long at both heights where it was evaluated (Figure 2B). Full and trimmed truth gave the same ordering of method means at each height. The *M*_*2*_ and *M*_*4*_ scores were closely similar. Across all 100 histories and three standard heights, replacing the classical NJ top of *M*_*4*_ by the anticentral top of *M*_*2*_ increased mean full-truth GRF by 0.00062 [0.00047, 0.00076] and trimmed-truth GRF by 0.00081 [0.00065, 0.00097]. The much larger gains over pooled NJ were shared by both biopsy-guided methods. The prespecified *M*_*2*_ − *M*_*0*_ comparison gave a mean within-history effect of 0.05727 [0.05358, 0.06108]. The improvement was present at every tested depth, although its magnitude decreased as the true tree grew. Topology-only MEDICC2 [14] provides a stronger contextual baseline than *M*_*0*_. The mean *M*_*2*_-minus-MEDICC2 effect was 0.02563 [0.02335, 0.02796]. These advantages persisted against trimmed truth: the three-height mean *M*_*2*_ − *M*_*0*_ and *M*_*2*_-minus-MEDICC2 effects were 0.07153 and 0.03425, respectively. Across the 300 standard conditions, *M*_*2*_ exceeded MEDICC2 in 275 full-truth and 280 trimmed-truth comparisons. The original primary comparison is reported in Table 4 and Figure 6. Table 13 gives the additional full/trimmed comparisons.

NestedBD-Long [17] exceeded pooled NJ in mean similarity at both evaluated heights. It also exceeded MEDICC2 at height 14, whereas MEDICC2 was stronger at height 24. Averaging both runs and both depths within each of the 98 complete histories, the mean *M*_*2*_-minus-NestedBD-Long effect was 0.02011 [0.01689, 0.02328] for full truth and 0.02702 [0.02312, 0.03088] for trimmed truth. The corresponding history-level wins/ties/losses were 86/2/10 and 88/2/8. The comparison describes trees obtained under the fixed inference budget. Scalar and between-run tree diagnostics did not establish MCMC convergence, particularly at height 24 (Supplementary Results 11.7).

### Closed-state recovery depends on depth and endpoint

The closed-state AD-F1 ranking changed with evolutionary depth (Figure 2C). Method *M*_*6*_ had the highest mean AD-F1 at height 14, exceeding pooled *M*_*5*_ by 0.08985. Pooled *M*_*5*_ had the highest mean at heights 24 and 34, with the *M*_*6*_ − *M*_*5*_ difference reaching −0.05116 at height 34. Similarly, *M*_*7*_ exceeded *M*_*3*_ at the two shallower depths, but their mean difference at height 34 was −0.00215, with an interval spanning zero. These prespecified descriptive AD-F1 comparisons show a depth-dependent ranking (Table 5). Rooted GRF showed a different pattern: *M*_*6*_ had the highest mean at every height, followed by *M*_*7*_. Both attained higher mean similarities than their pooled counterparts, although all five methods’ mean similarities decreased with depth (Table 6). The same ordering of mean GRF scores and decline with depth held against trimmed truth (Supplementary Figure 7C). The effects of standard, late, and random sampling varied across methods and endpoints (Table 10).

### Pipeline resources

Including condition-specific construction of *D*, the mean *M*_*2*_ pipeline runtime was 2.03 seconds, compared with 10.33 seconds for MEDICC2. Maximum resident memory was 172 MB for *M*_*2*_ and 453 MB for MEDICC2. Both measurements used four concurrent case workers and excluded evaluation (Table 11). Thus *M*_*2*_ combined higher partial-tree similarity with lower measured pipeline time and memory on these inputs. Complete scores, paired contrasts, and resource accounting are given in Supplementary Results 11.

## 6 Empirical feasibility on longitudinal breast-cancer data

To complement the truth-based simulations, we assessed genome-scale feasibility using public single-nucleus DNA-sequencing data for KTN152, patient P14 in the longitudinal triple-negative breast cancer study of Kim et al. [16]. Tumor specimens were collected before neoadjuvant chemotherapy, after two treatment cycles, and at surgery after six cycles. We aligned the single-end reads to hg19 and jointly processed all biopsies with `CopyKit` [26]. Biopsy labels were excluded from segmentation, integer calling, nearest-neighbor smoothing, and quality control, which all preceded reconstruction. Complete processing and provenance are reported in Supplementary Methods 12.

Absolute copy-number scaling remained unresolved. The available KTN152 data lacked numerical DNA-content measurements, and a reconstruction-blind comparison with `scAbsolute` [23] yielded discordant estimates for these amplified single-end libraries. We therefore fixed an exploratory feasibility analysis before reconstruction, using the *k* = 4 smoothed `CopyKit` estimates that passed prespecified technical and profile-correlation criteria. These estimates were retained without subsequent rescaling. Biopsy-specific filtering produced an uneven retained sample, whose counts describe the analyzed data rather than clone prevalences. The inferred trees are evaluated descriptively because an empirical reference topology is unavailable.

The frozen input contained 79 cells, distributed as 28, 16, and 35 across the successive biopsies. Within-biopsy collapse yielded 77 biopsy-indexed occurrences and 76 distinct states, retaining separate temporal occurrences of the one recurrent state (Figure 3A). Each profile comprised 12,167 variable-width bins on chromosomes 1–22 and X (Figure 3B). Of 936,859 occurrence–bin entries, 5,669 (0.61%) had copy number at least 8. Among these, 2,761 (0.29% of all entries) exceeded the pinned MEDICC2 input cap of 8. The maximum estimate was 40. We combined independently computed chromosome-specific `cnp2cnp` costs into one symmetric whole-genome matrix and supplied it unchanged to *M*_*0*_ and *M*_*2*_.

**Figure 3.**
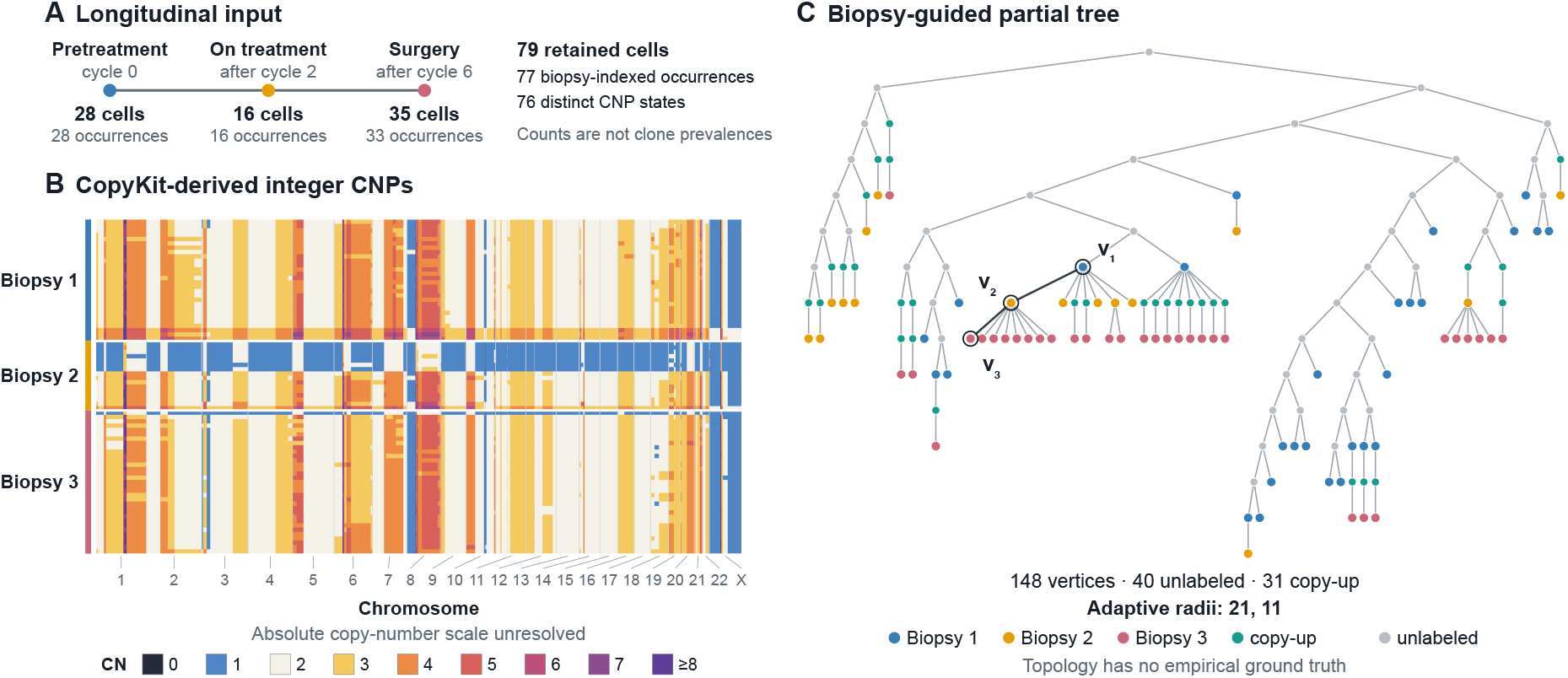
Empirical feasibility on longitudinal KTN152 CNP estimates. Biopsies 1–3 are numbered chronologically and correspond to *B*_*0*_, *B*_*1*_, *B*_*2*_, respectively. **(A)** Retained cells and within-biopsy-collapsed occurrences from the three successive biopsies. Counts reflect post-filtering sample sizes. **(B)** CopyKit-derived integer profiles grouped by biopsy and ordered within each biopsy by mean copy number and then state identifier, independently of either reconstruction. Values of at least eight share one display color. Chromosome boundaries follow bin coordinates, with leader lines separating labels for the shorter chromosomes. The absolute copy-number scale remains unresolved. **(C)** The *M*_*2*_ partial CNP-tree. Blue, orange, and rose vertices are observed biopsy occurrences. Teal vertices are inferred copy-up occurrences, and grey vertices are unlabeled top vertices. The two biopsy transitions used adaptive radii 21 and 11. The highlighted path *v*_*1*_ → *v*_*2*_ → *v*_*3*_ joins observed occurrences from the first, second, and third biopsies, respectively. Vertical position denotes graph depth. The topology is an exploratory reconstruction, with no empirical reference tree available for validation.

Both methods returned valid partial CNP-trees containing all 76 states. The *M*_*0*_ tree had 151 vertices, 75 unlabeled vertices, and maximum depth 18. The *M*_*2*_ tree had 148 vertices: 77 observed occurrences, 31 inferred copy-up occurrences, and 40 unlabeled top vertices, with maximum depth 14 (Figure 3C). Biopsy colors distinguish observed occurrences, including the two observations of the recurrent state. The highlighted path *v*_*1*_ → *v*_*2*_ → *v*_*3*_ links a pretreatment occurrence to an on-treatment occurrence and a surgical occurrence. It illustrates model-inferred CNP-state ancestry across the three biopsies. Across the two biopsy transitions, *M*_*2*_ selected radii 21 and 11, made 31 copy-up decisions, and deferred six minimum-distance ties. Matrix construction took 4.20 seconds with six workers, and the two reconstructions took 1.69 and 1.73 seconds (Table 16). Together, these outputs demonstrate computational feasibility on longitudinal, patient-derived CNP estimates at whole-genome resolution.

## 7 Discussion

We formulated partial and closed-state CNP-tree reconstruction for longitudinal biopsies in a model that distinguishes evolutionary occurrences from copy-number states and permits the same observed state to recur. The biopsy-guided construction separates temporally supported connections between successive samples from completion of the unresolved ancestral top. This decomposition accommodates both partial outputs with latent internal nodes and closed-state outputs that reuse observed CNPs. Given *D*, the eight evaluated CTBF methods run in polynomial time under unit-cost arithmetic.

In the prospectively specified independent simulation study, the biopsy-guided partial method improved similarity over pooled `cnp2cnp`–NJ in every history block. The partial-tree comparisons support this pattern under both full and trimmed truth. Methods *M*_*2*_ and *M*_*4*_ exceeded topology-only MEDICC2 [14] in mean similarity at all three heights and finite-budget NestedBD-Long at heights 14 and 24. NestedBD-Long exceeded MEDICC2 at height 14, whereas their order reversed at height 24. The two biopsy-guided partial methods gave closely similar scores. Their shared bottom construction accounts for most of the observed gain over pooled NJ, with a small additional gain from anticentral top completion (Supplementary Results 11.5). Closed-state AD-F1 rankings changed with depth, whereas *M*_*6*_ had the highest mean rooted-GRF similarity at each height. Additional biopsies improved mean AD-F1 for both biopsy-guided closed-state methods when scoring all observed states in each condition. The fixed-label control also showed improved ancestry recovery, with larger gains than for the pooled counterparts using the same completion rule. The pooled anticentral method also benefited from additional profiles. Together, these results support the biopsy-guided construction under the tested CTBF data-generating model. Method selection must nevertheless reflect the output class, evolutionary regime, and evaluation endpoint.

### Scope and limitations

The simulation study used exact CNPs from genotype-state histories, with whole-genome doubling disabled. Clone abundance, spatial sampling, sequencing noise and copy-number calling error lie outside this setting. Total-CNP observations also leave copy-neutral loss of heterozygosity and copy-number-preserving rearrangements unresolved. For KTN152, uncertainty in absolute copy number and the absence of a known topology restrict the empirical assessment to computational feasibility on patient-derived estimates.

The external comparisons assess rooted topology. NestedBD-Long models a sampled birth– death process, whereas the simulator retains genotype-state representatives. Its accuracy summaries describe finite-budget trees from inputs with two successful runs. The failed runs and inconclusive convergence diagnostics are reported in Supplementary Methods 11.7. Both GRF targets retain recurrent-label multiplicities and therefore assess the specified occurrence-tree representation.

These findings motivate further evaluation under alternative evolutionary and observation models, longitudinal data with externally validated absolute copy number, and extensions that infer ancestral CNPs beyond the observed profile set.

## Code and data availability

CTBF 1.2.0 is available at https://github.com/j-paszek/ctbf. The release includes simulation, reconstruction, and evaluation software, experimental configurations, and the scripts and numerical evidence used to reproduce the simulation tables and figures. The source snapshot is archived on Zenodo at https://doi.org/10.5281/zenodo.23366634.

The companion ctbfbio repository is available at https://github.com/j-paszek/ctbfbio. Its release assets provide the KTN152 preprocessing scripts, integer CNP matrices, genomic-bin coordinates, biopsy and cell mappings, and empirical analysis summaries. The repository snapshot for release 1.0.1 is archived on Zenodo at https://doi.org/10.5281/zenodo.23369066. The original sequencing reads are available from the European Nucleotide Archive under accession PRJNA396019 (SRA study SRP114962).

## Funding

This work was supported by the National Science Centre under grant no. 2020/39/D/ST6/03321.

## 8 Supplementary formal context

### 8.1 Tree levels and prefixes

For an integer *i* ≥ 0, the *ith level* of *T* is the set of vertices at depth *i*. A profile assignment or labeling on *T* is restricted to the same vertex set when a tree prefix is taken. When a prefix is treated as an evolutionary history tree, its profile set is the image of the restricted profile assignment, and its state labels and profile-to-label bijection are inherited by restriction.

### 8.2 Relations to established tree formalisms

Our CNP-tree definition is related to the copy-number-tree model of El-Kebir et al. [9]. Both formalisms assign a CNP to every node and permit the same profile at distinct nodes. Their optimization problem additionally requires a full binary topology with a diploid root and observed profiles at the leaves, assigns a directed event sequence to every edge, and minimizes total event cost. Our definition permits a nonbinary topology with any root profile in *P* and allows profiles in *P* at leaves or internal nodes. It specifies the tree representation independently of edge-event labels and optimization criteria.

Partial CNP-trees are related to the semi-labeled trees of Deng and Fernández-Baca [6]. Both require every leaf to be labeled and allow internal nodes to be labeled. In their formalism, each taxon label maps to one node, several taxon labels may map to the same node, and every node of degree at most two must carry a label. In our formalism, each labeled node maps to one CNP-state label, the same state label may occur at distinct nodes, and only leaves are required to be labeled. Multiset-labeled tree formalisms likewise permit repeated labels [15].

The copied occurrences used below differ from placing an observed taxon or cell itself at an ancestral vertex, as in generally labeled Family-Joining trees and MEDALT spanning trees [13, 25]. LACE 2.0 instead constructs a longitudinal representation of an inferred clone tree by propagating clones across sampling times. It can insert an earlier occurrence of a parental clone when that clone is first observed later than its child [1]. Our bottom construction copies the child’s state when a unique plausible parent cannot be selected, retaining this unresolved assignment for the next earlier transition. The temporal construction is related to SeqTrack, which represents sampled isolates by individual nodes and infers an optimum branching from strictly earlier to later samples [12]. Our temporal method admits same-biopsy ancestry and adds a directional copy-number plausibility criterion.

### 8.3 Non-identifiability of ancestry

#### Proposition 3.

*Fix four distinct CNP-state labels a, b, c, d, take B*_*0*_ = *{b, c} and B*_*1*_ = *{d, a} at times τ*_*0*_ *< τ*_*1*_, *and fix a common profile map ξ and an admissible dissimilarity matrix D on U* = *{a, b, c, d}. There exist two rooted, state-labeled true trees with the same five occurrences, the same state assignment, and the same sampling map that are both compatible with the identical input ℐ* = (*B, ξ, D*) *and satisfy:*

a. *the sampled-occurrence ancestry differs. In one tree v*_*b*_ *is an ancestor of v*_*d*_, *whereas in the other v*_*c*_ *is an ancestor of v*_*d*_. *Consequently, the input does not identify one true sampled ancestry*.
b. *in both trees the unobserved root r carries state a and is an ancestor of the sampled occurrences v*_*b*_ *and v*_*c*_ *from B*_*0*_, *while the distinct occurrence v*_*a*_ *carrying the same state is sampled only in B*_*1*_.

*Proof*. Fix four distinct CNPs with state labels *a, b, c, d*, and take the two biopsies *B*_*0*_ = *{b, c}* and *B*_*1*_ = *{d, a}* at times *τ*_*0*_ *< τ*_*1*_. Let *V* = *{r, v*_*a*_, *v*_*b*_, *v*_*c*_, *v*_*d*_*}*, let *r* be the root, assign state *a* to *r*, and assign state *s* to *v*_*s*_ for *s* ∈ *{a, b, c, d}*. Define

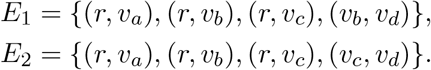

For both trees use

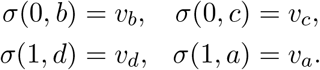

The maps preserve states and satisfy the biopsy-order constraint. The profile map, biopsy times, and dissimilarity matrix are identical for both histories, so they induce the same input *ℐ*. In *E*_*1*_, *v*_*b*_ is an ancestor of *v*_*d*_ and *v*_*c*_ is not. In *E*_*2*_, *v*_*c*_ is an ancestor of *v*_*d*_ and *v*_*b*_ is not. Thus the two compatible histories have different sampled ancestry, while the root *r* is unobserved, carries state *a*, and is ancestral to both *v*_*b*_ and *v*_*c*_. The state *a* is first sampled in *B*_*1*_ at the distinct occurrence *v*_*a*_.

Figure 4 illustrates these two histories. Part (a) is an identifiability limitation. A reconstruction selected from *ℐ* cannot be guaranteed to coincide with the unknown true sampled ancestry for every compatible history, so an ideal score is not guaranteed even when the reconstruction is feasible.

**Figure 4.**
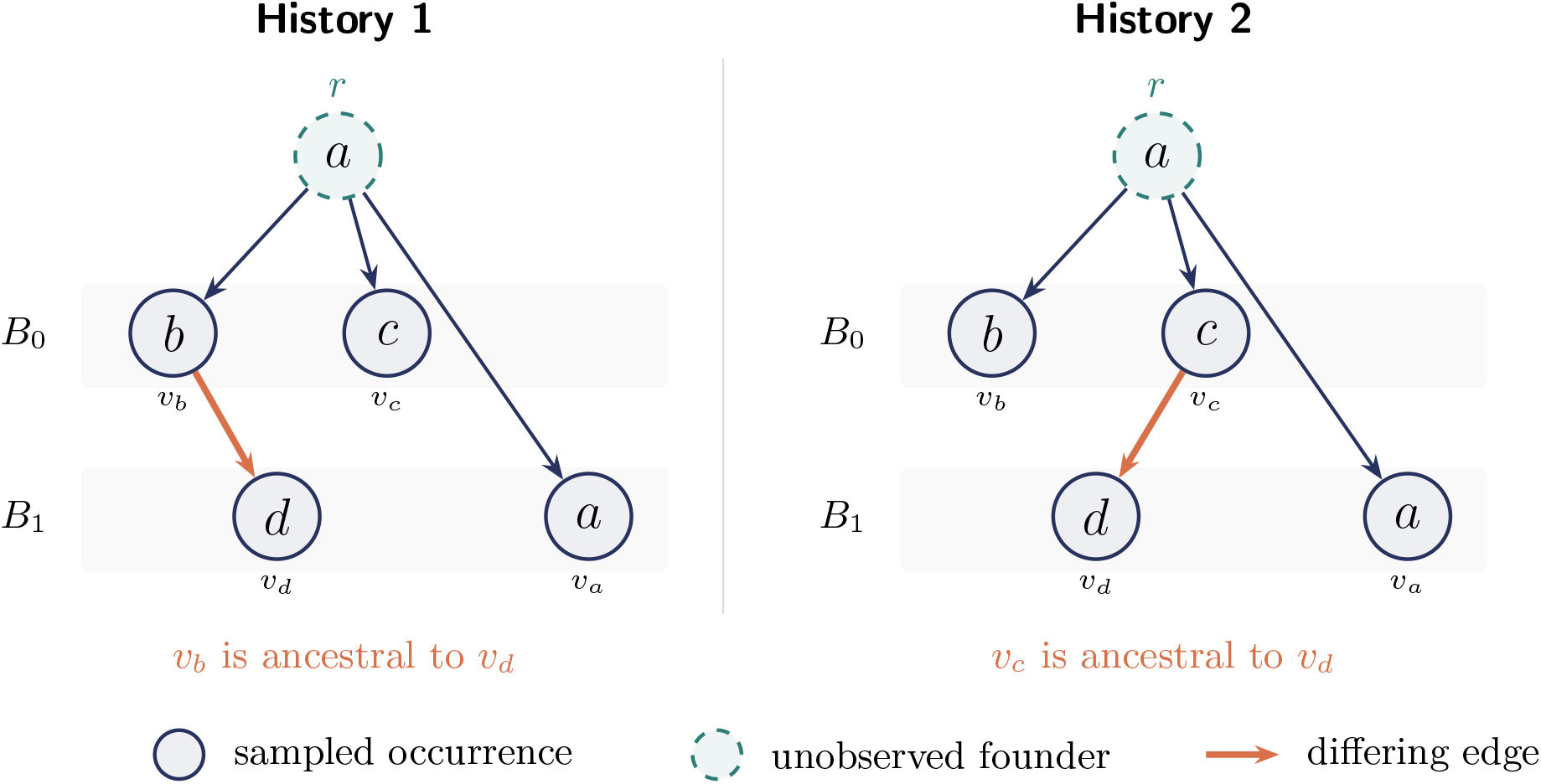
Two true histories compatible with the same input *ℐ*: *B*_*0*_ = *{b, c}, B*_*1*_ = *{d, a}, τ*_*0*_ *< τ*_*1*_, and identical profile map *ξ* and dissimilarity matrix *D*. Both have five occurrences. Node text gives the CNP state, and shaded rows identify sampled occurrences. The highlighted edge makes *v*_*d*_ a descendant of *v*_*b*_ in History 1 and of *v*_*c*_ in History 2. The unobserved founder *r* and the later sampled occurrence *v*_*a*_ are distinct nodes carrying state *a*.

**Figure 5.**
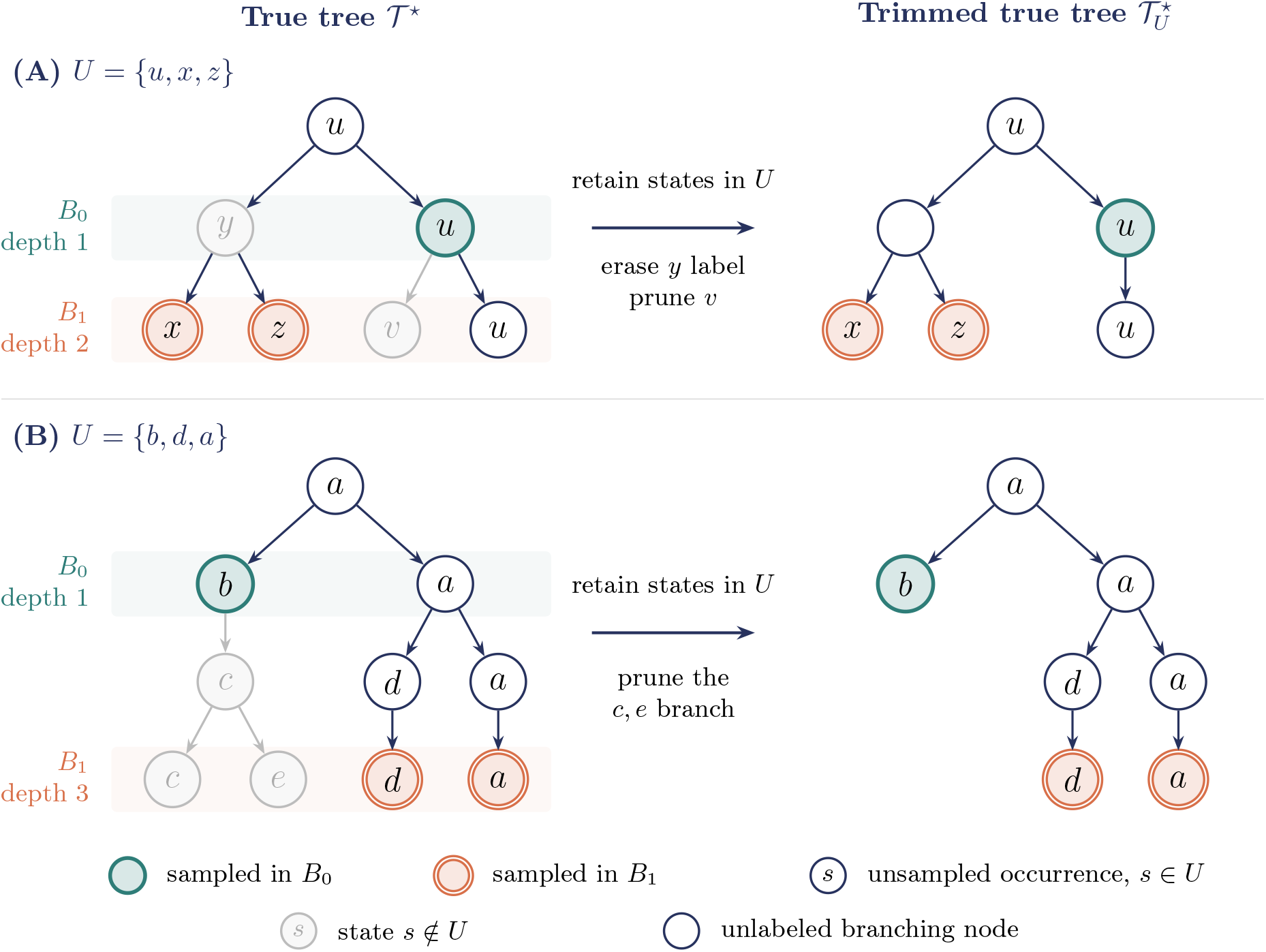
State-restricted true trees. Letters denote CNP-state labels. Colored nodes identify sampled occurrences, with *B*_*0*_ in teal and *B*_*1*_ in orange. Biopsy depths are measured from the original root at depth zero. (A) For *B*_0_ = *{u}* and *B*_1_ = *{x, z}*, the *y* label is removed but its branching node remains, whereas the *v* leaf is pruned. All three occurrences of *u* survive, although only the middle one was sampled. (B) For *B*_0_ = *{b}* and *B*_1_ = *{d, a}*, the branch below *b* is pruned. All *a* and *d* occurrences survive, including the labeled unary paths. An empty circle denotes an unlabeled node, not a new state.

Part (b) concerns the same histories. Temporal compatibility constrains relations between sampled occurrences, not the first biopsy in which a state label appears. Therefore a state observed in *B*_*1*_ is not excluded from labeling an unobserved occurrence on the path to an occurrence sampled in *B*_*0*_. The deferred copy-up operation preserves this possibility by creating an earlier occurrence with the child’s state when no unique plausible parent is available. It can therefore represent the unobserved state-*a* founder shown in Figure 4.

## 9 Supplementary reconstruction algorithms

### 9.1 Operational conventions

Whenever a reconstruction method assigns an edge weight, the weight is an auxiliary output attribute and does not change the topology or node-labeling class. Every randomized rule uses a pseudorandom generator initialized by the prespecified method seed.

### 9.2 Secondary method compositions

The principal comparison includes eight CTBF methods and the external comparators MEDICC2 and NestedBD-Long. Table 2 identifies the two additional global occurrence-level constructions.

**Table 2.**
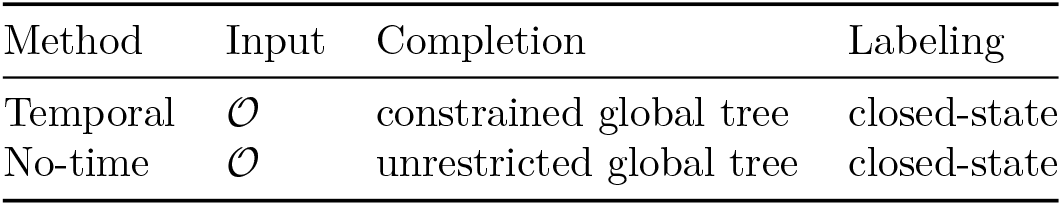
Secondary reconstruction-method compositions.

**Table 3.** Standard synthetic evaluation regime. Probabilities are conditional as stated. Resource limits are operational ceilings rather than biological parameters.

| Component | Standard value or rule |
| --- | --- |
| Genome and founder | $m = 100$ bins on one synthetic chromosome. Initial copy number is 2 in every bin. |
| Evolutionary depth | $H \in \{14, 24, 34\}$. The standard, low-depth, and stress conditions use $H = 24$, $H = 14$, and $H = 34$, respectively. |
| Descendant generation | One baseline and one additional descendant attempt per retained occurrence. No fitness or state-lineage regulation. |
| Segmental initiation | Constant schedule with per-bin probability $e = 0.1/m = 0.001$, hence 0.1 expected initiation proposals per attempted descendant. |
| Direction and footprint | The conditional gain probability is 0.5. The conditional interval probability is 0.1, with a singleton event otherwise. Interval direction is sampled left or right with equal probability, and the endpoint is uniform within the same chromosome. Singleton intervals are retained. |
| Magnitude | Singleton events and interval losses change copy number by one. Conditional on an interval gain, +1 has probability 0.8 and +2 has probability 0.2. |
| Viability and retention | Copy number zero is absorbing. Descendants with an all-zero profile, or with copy number zero in any of ten prospectively fixed crucial bins, are rejected. At most one occurrence per exact CNP is retained within a generation. |
| Disabled mechanisms | Whole-genome doubling, telomeric enrichment, multiplicative gains, and soft state-lineage regulation are disabled. |
| Resource guard | At most 2,000 retained genotype-state occurrences per generation and 40,000 total true-tree nodes. Exceeding either limit yields a recorded failed replicate without truncation or replacement. |

**Table 4.** Partial-tree confirmation. Panel (a) gives absolute mean full-truth rooted-GRF similarity for the four methods evaluated at all three heights. Panel (b) gives paired effects with the first method favored by positive values. Height rows contain 100 condition pairs. Block-mean rows contain 100 independent effects, each averaged over the three heights.

**(a) Absolute mean rooted GRF**
| $H$ | $M_0$ | MEDICC2 | $M_2$ | $M_4$ |
| --- | --- | --- | --- | --- |
| 14 | 0.188083 | 0.246998 | 0.285265 | 0.284549 |
| 24 | 0.111363 | 0.133827 | 0.154148 | 0.153401 |
| 34 | 0.089043 | 0.102587 | 0.120890 | 0.120494 |

**(b) Paired rooted-GRF effects**
| Contrast | $H$ | Mean effect | 95% bootstrap interval | W/T/L |
| --- | --- | --- | --- | --- |
| $M_2 - M_0$ | 14 | 0.097182 | 0.087487–0.107013 | 99/0/1 |
|  | 24 | 0.042785 | 0.038345–0.047540 | 100/0/0 |
|  | 34 | 0.031848 | 0.029646–0.034024 | 99/0/1 |
|  | block mean | 0.057271 | 0.053579–0.061079 | 100/0/0 |
| $M_2 - \text{MEDICC2}$ | 14 | 0.038267 | 0.032027–0.044560 | 91/0/9 |
|  | 24 | 0.020321 | 0.016725–0.023964 | 89/0/11 |
|  | 34 | 0.018304 | 0.015989–0.020562 | 95/0/5 |
|  | block mean | 0.025630 | 0.023354–0.027962 | 100/0/0 |

**Table 5.** Closed-state confirmation under the three standard biopsy conditions. Panel (a) gives mean AD-F1. Panel (b) gives the two prespecified biopsy-guided-minus-pooled paired contrasts.

| (a) Absolute mean AD-F1 |  |  |  |  |  |
| --- | --- | --- | --- | --- | --- |
| $H$ | $M_1$ | $M_3$ | $M_5$ | $M_6$ | $M_7$ |
| 14 | 0.446200 | 0.517410 | 0.788337 | 0.878188 | 0.791407 |
| 24 | 0.454830 | 0.615504 | 0.771636 | 0.756390 | 0.702623 |
| 34 | 0.439051 | 0.663548 | 0.687091 | 0.635930 | 0.661399 |

| (b) Paired AD-F1 effects |  |  |  |  |
| --- | --- | --- | --- | --- |
| Contrast | $H$ | Mean effect | 95% bootstrap interval | W/T/L |
| $M_6 - M_5$ | 14 | 0.089851 | 0.050772–0.134667 | 63/8/29 |
|  | 24 | −0.015246 | −0.040537–0.015386 | 35/4/61 |
|  | 34 | −0.051161 | −0.069427–−0.034131 | 29/0/71 |
|  | block mean | 0.007815 | −0.010314–0.028785 | 44/0/56 |
| $M_7 - M_3$ | 14 | 0.273997 | 0.223112–0.328521 | 94/0/6 |
|  | 24 | 0.087118 | 0.054496–0.123088 | 73/0/27 |
|  | 34 | −0.002149 | −0.021285–0.017280 | 44/0/56 |
|  | block mean | 0.119656 | 0.093285–0.148428 | 85/0/15 |

**Table 6.** Complementary closed-state rooted-GRF similarity on the three standard conditions. Panel (a) gives mean scores over 100 histories at each height. Panel (b) gives paired differences with two-sided 95% history-block bootstrap intervals and wins/ties/losses. A block-mean difference averages the three heights within one history. Panels (a) and (b) use full truth. Panel (c) gives mean scores against trimmed truth for the same five methods and histories. All rows are descriptive and retain 100 independent history blocks.

| (a) Absolute mean full-truth rooted GRF |  |  |  |  |  |  |
| --- | --- | --- | --- | --- | --- | --- |
| $H$ | $M_1$ | $M_3$ | $M_5$ | $M_6$ | $M_7$ | |
| 14 | 0.206792 | 0.142966 | 0.212162 | 0.287347 | 0.273672 |  |
| 24 | 0.120426 | 0.069558 | 0.121825 | 0.155008 | 0.146854 |  |
| 34 | 0.094387 | 0.049816 | 0.095742 | 0.121448 | 0.117007 |  |

| (b) Paired full-truth rooted-GRF effects |  |  |  |  |
| --- | --- | --- | --- | --- |
| Contrast | $H$ | Mean effect | 95% bootstrap interval | W/T/L |
| $M_6 - M_5$ | 14 | 0.075185 | 0.065290–0.085118 | 96/0/4 |
|  | 24 | 0.033182 | 0.028556–0.038098 | 94/0/6 |
|  | 34 | 0.025705 | 0.022951–0.028331 | 96/0/4 |
|  | block mean | 0.044691 | 0.041169–0.048355 | 100/0/0 |
| $M_7 - M_3$ | 14 | 0.130706 | 0.122436–0.138825 | 100/0/0 |
|  | 24 | 0.077297 | 0.073857–0.080931 | 100/0/0 |
|  | 34 | 0.067191 | 0.065260–0.069083 | 100/0/0 |
|  | block mean | 0.091731 | 0.088795–0.094707 | 100/0/0 |

| (c) Absolute mean trimmed-truth rooted GRF |  |  |  |  |  |
| --- | --- | --- | --- | --- | --- |
| $H$ | $M_1$ | $M_3$ | $M_5$ | $M_6$ | $M_7$ |
| 14 | 0.217871 | 0.150314 | 0.223502 | 0.308486 | 0.292365 |
| 24 | 0.145176 | 0.084206 | 0.147191 | 0.192974 | 0.182731 |
| 34 | 0.126038 | 0.067354 | 0.127962 | 0.169081 | 0.163272 |

**Table 7.**
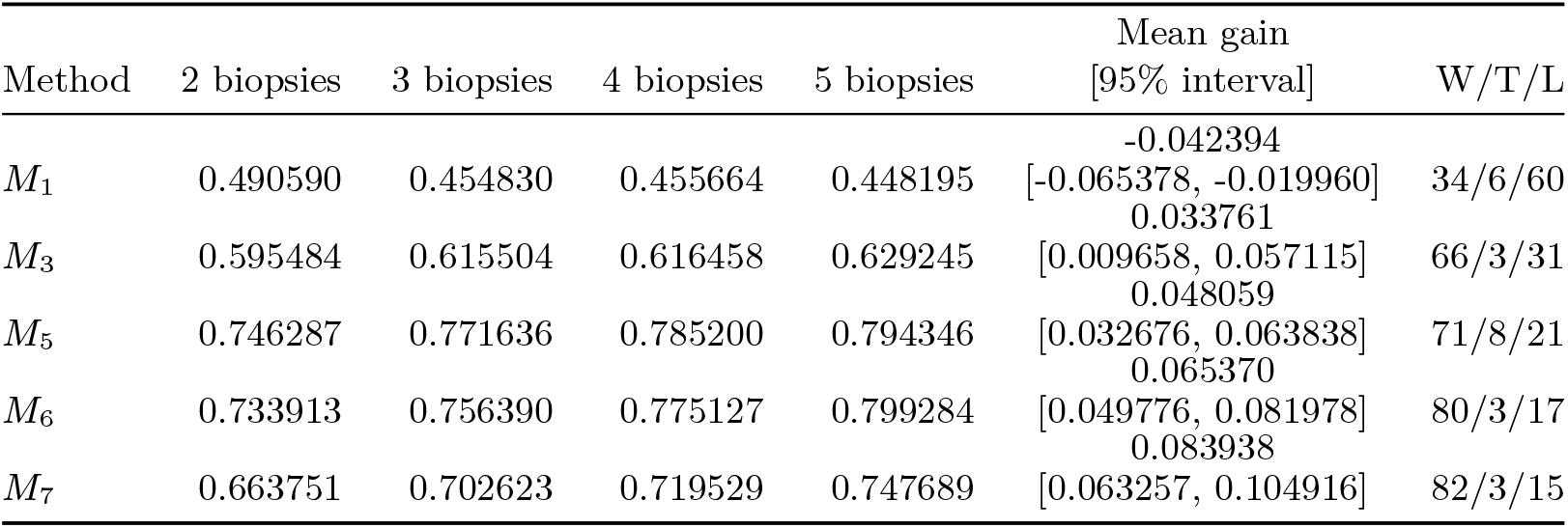
Ordinary closed-state AD-F1 by biopsy number, using all observed states of each condition. Means and five-minus-two differences use all 100 histories. Intervals are pointwise 95% paired-history bootstrap intervals. W/T/L counts history-level changes.

### 9.3 Adaptive/deferred bottom construction

Use the level-extended sets 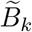, initialized forest *F*_*B*_, radii *r*_*t*_, and transition sets *P*_*t*−*1*_ and *Y*_*t*_ defined in Section 3.2.1. For each initialized occurrence *v*_*k,s*_, set *κ*(*v*_*k,s*_) = *k* and *ℓ*_*B*_(*v*_*k,s*_) = *s*. For *s* ∈ *B*_*k*_, set *η*(*k, s*) = *v*_*k,s*_. The nearest-rank convention selects the lower of the two central values when 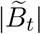 is even.

At transition *t* − 1 → *t*, for *y* ∈ *Y*_*t*_, write *s* = *ℓ*_*B*_(*y*) and define

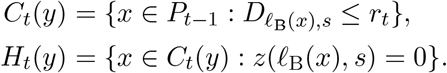

If 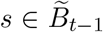, then *v*_*t*−*1,s*_ exists and belongs to *C*_*t*_(*y*) because *D*_*ss*_ = 0 ≤ *r*_*t*_. Select it as parent. Otherwise, select a parent exactly when the minimum of 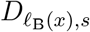 over *H*_*t*_(*y*) is attained by one candidate. The decision is deferred when *H*_*t*_(*y*) is empty or the minimum is tied. Hence candidate enumeration never resolves an exact distance tie.

Every selected parent *x* receives an edge *x* → *y* of weight 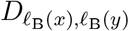. Several children may attach to one parent. Once all children have been examined against the unchanged snapshot *P*_*t*−*1*_, create for each deferred *y* a fresh occurrence 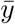 at index *t* − 1, set 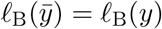, and add 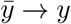 with weight zero. These copies are inserted as one batch. Thus no copy can parent another child in the transition that created it, but it participates in *Y*_*t*−*1*_ during the next transition. The child and its copy remain distinct evolutionary occurrences carrying the same state.

#### Proposition 4

(Feasibility of bottom–top reconstruction). *For every reconstruction input ℐ, the adaptive/deferred construction terminates with a biopsy-guided bottom forest and an injective observation map η. Every partial top completion of this forest is a feasible partial reconstruction. Every closed-state top completion is a feasible closed-state reconstruction, with the same observation map*.

*Proof*. Initialization creates a distinct occurrence *v*_*k,s*_ for every 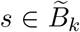. Thus *η*(*k, s*) = *v*_*k,s*_ is injective on *O* and preserves both state labels and biopsy indices. At transition *t* − 1 → *t*, every current root at index *t* receives exactly one parent at index *t* − 1, either from the frozen candidate set or as a fresh same-state copy. No other vertex receives an incoming edge in that transition. Every added edge strictly increases *κ*, so no directed cycle can arise. The graph remains a rooted forest. After the *q* − 1 transitions, every positive-index vertex has a parent, whereas all index-0 vertices remain roots. All original occurrences and labels are retained, and the resulting forest satisfies the bottom-forest requirements.

A top completion preserves this forest and attaches new edges only at component roots and introduced top nodes. All leaves remain labeled because every introduced top node is ancestral to a forest root. Within an original component, ancestry is unchanged and every directed edge increases *κ*. A directed path between observations in different original components can leave the source component only at its root. Its source observation must therefore have biopsy index 0, which cannot exceed the target index. Consequently, 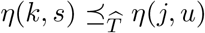 implies *j* ≤ *k*. The observation map remains injective and label-preserving, and every state in *U* remains represented. Leaving new top nodes unlabeled or labeling them by states in *U* gives the respective feasible output classes.

### 9.4 Binary anticentral top completion

We use the pair score *S*_*ij*_ and diploid deviation *ϕ* defined in Section 3.4. The initialization, tie conventions, and updates are specified below. Let *ℱ* = (*F, ℓ*_*F*_) have an ordered list of *k*_*0*_ component roots (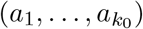). Denote the operational labeling used during completion by *ℓ*. Initially, *ℓ* = *ℓ*_*F*_ on *V*_*F*_, and *ℓ* is extended whenever a copied occurrence is introduced. Initialize

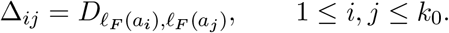

When *k*_*0*_ *>* 1, define

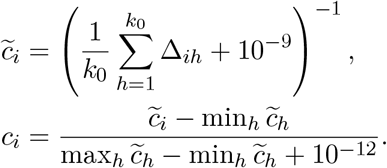

Thus *c*_*i*_ ∈ [0, 1], with larger values representing smaller initial mean dissimilarity.

At a reduction with *k* ≥ 2 active roots, set

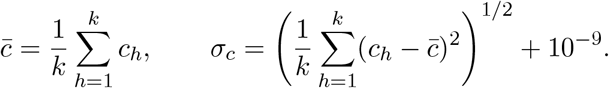

For every *i < j*, independently draw *Z*_*ij*_ uniformly from [0, 1) using the fixed seeded generator and compute the pair score *S*_*ij*_. Select the minimum-scoring pair. The perturbation is applied to every pair score before selection. A residual numerical tie is resolved by the fixed active-root order.

For selected roots *a*_*i*_, *a*_*j*_, the only zero-regain-plausible direction is chosen when exactly one exists. If both are plausible, choose the root with smaller *ϕ*. If their deviations are equal, or neither direction is plausible, choose the root with larger current *c*. An exact centrality tie is resolved by the seeded generator. Denote the resulting parent and child indices by *i*_*p*_ and *i*_*c*_.

Create a fresh active occurrence *a*_*new*_ with 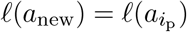 and add 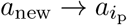 and 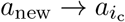, both with weight 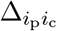. Replace the active parent by *a*_*new*_, remove the child, retain the parent dissimilarity row,

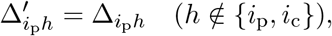

and update

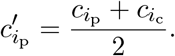

All other retained dissimilarities and centralities remain unchanged. Continue until one active root remains. Every reduction introduces a two-child parent, so the added top skeleton is binary. A one-component forest is returned unchanged. Suppressing the labels of introduced top nodes gives the partial projection, whereas retaining their copied parent-state labels gives the closed-state projection.

### 9.5 Classical neighbor-joining partial completion

Cordonnier and Lafond reconstructed unrooted, leaf-labeled trees by applying PHYLIP NJ to CNP dissimilarities [4]. Applied to *ℱ*_*U*_, the CTBF procedure below is the cnp2cnp–NJ baseline *M*_*0*_. It implements the classical NJ criterion within CTBF but returns the rooted representation required by the reconstruction interface. Applied to the biopsy-guided bottom forest *ℱ*_*B*_, the same completion defines the partial method *M*_*4*_. A root must be specified to represent the original unrooted output as a partial CNP-tree. The published experiment used CNPs from true-tree leaves. Our pooled set *U* may also contain a state observed at an ancestral occurrence, but *M*_*0*_ nevertheless represents every state as a leaf. Method *M*_*0*_ is therefore a pooled state-level distance-tree baseline.

Let *ℱ* = (*F, ℓ*_*F*_) have *k*_*0*_ component roots in the fixed active order 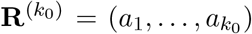. Initialize

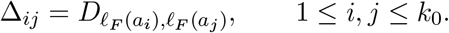

With *k >* 2 active roots, classical neighbor joining [21] computes

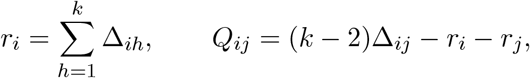

and selects a pair *i < j* minimizing *Q*_*ij*_. An exact tie is resolved by the first pair in the current active order. For the selected pair, define

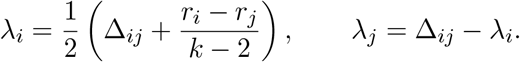

The reduction introduces an unlabeled node *u*, adds the edges *u* → *a*_*i*_ and *u* → *a*_*j*_ with weights *λ*_*i*_ and *λ*_*j*_, and replaces the selected roots by *u*. For every other active root *a*_*h*_, the updated dissimilarity is

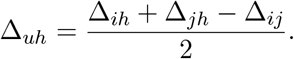

Unselected roots retain their relative order and *u* is appended to the active list. Negative limb lengths are retained.

When two active roots remain, an unlabeled synthetic root is connected to both with half their remaining dissimilarity. This final operation gives a rooted representation under the final-join rooting convention. If *k*_*0*_ = 1, the sole component root is returned unchanged. The implementation uses 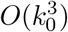 arithmetic operations and 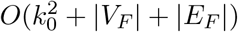 memory, excluding the stored CNP profiles. All introduced nodes remain unlabeled. Consequently, *M*_*0*_ returns a partial CNP-tree without inferring ancestral CNP states. It need not satisfy the observation-map requirements of the longitudinal reconstruction problem.

### 9.6 MEDICC2 topology-only comparator

MEDICC2 obtains a symmetric minimum-event distance by minimizing two directed evolutionary paths through a possible common ancestor. It infers copy-number phylogenies from these distances [14]. Its standard workflow applies NJ to the pairwise distances and can subsequently reconstruct ancestral copy-number profiles. We used the topology stage, retaining unlabeled internal nodes. The pair-specific ancestors used in distance calculation remain implicit.

Every observation (*k, s*) ∈ *O* was exported as a distinct opaque taxon. The exported table contained neither biopsy indices nor original state labels. Equal profiles observed at different times were not deduplicated. The input comprised integer total copy numbers on the common 100-bin chromosome together with an artificial diploid normal. We used MEDICC2 version 1.1.2 from the official source revision `3f1b8a6` with options –total-copy-numbers, `–topology-only, –no-wgd, –maxcn 8`, and `–n-cores 1`. Disabling whole-genome doubling matches the standard synthetic regime and leaves a deliberate model mismatch for interval gains of magnitude two, which MEDICC2 represents by successive unit gains.

The returned PhyloXML tree was required to have one outer root whose children were the diploid-normal leaf and one tumor subtree. We removed the normal and the outer root, promoted the tumor-subtree root, mapped every remaining leaf back to its observation state, and left every internal node unlabeled. This truth-independent projection retains one leaf per member of *O* and produces a partial CNP-tree. It assigns no ancestral CNP and is therefore excluded from the closed-state AD-F1 comparison. Because MEDICC2 computes its own distances as well as its NJ topology, its comparison with *M*_*0*_ and *M*_*2*_ is a whole-pipeline comparison rather than a reconstruction-only comparison on the common matrix *D*. The MEDICC2 comparison comprised the 300 standard conditions. Biopsy-number and placement sensitivity were evaluated with all eight CTBF methods.

### 9.7 Labeled closest-pair completion

Let *ℱ* = (*F, ℓ*_*F*_) have ordered component roots 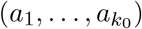. Denote the operational labeling by *ℓ*, initialize *ℓ* = *ℓ*_*F*_ on *V*_*F*_, and set 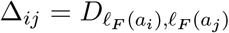. The labeling *ℓ* is extended whenever a copied occurrence is introduced. At each reduction, the labeled closest-pair completion selects

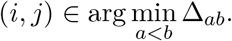

The first pair in active order is used when the minimum is attained more than once. Within the selected pair, the parent index is selected from

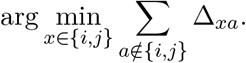

An exact parent tie is resolved equiprobably by the fixed seeded generator. Writing *i*_*p*_ and *i*_*c*_ for the parent and child indices, the reduction creates a fresh occurrence *u* with 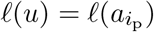. It adds 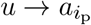 with weight 0 and 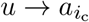 with weight 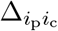. The active parent root is replaced by *u*, the child row and column are deleted, and every retained dissimilarity in the parent row remains unchanged. Repeating the reduction returns a rooted binary closed-state completion. A one-component forest is returned unchanged. The procedure requires 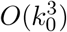 arithmetic operations and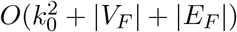 memory, excluding stored profiles.

### 9.8 Rooted labeled-*Q* completion

The rooted labeled-*Q* completion uses the neighbor-joining pair score [21] but contracts components without constructing an NJ latent node. Let *ℱ* = (*F, ℓ*_*F*_) have an initial ordered root list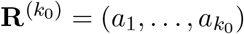. Define the fixed orientation score

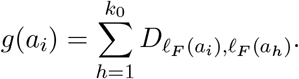

At a reduction with active list 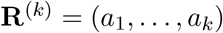 and *k >* 2, define

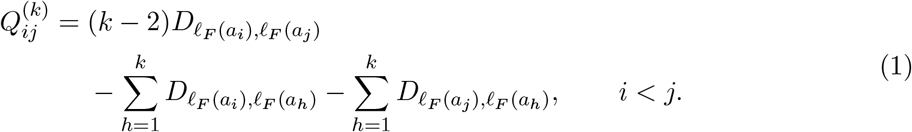

The ordered active-root list is algorithmic state. It determines the enumeration of exact pair ties but does not impose an evolutionary order on the roots. Algorithm S1 states the complete procedure.

**Algorithm S1: rooted labeled-***Q* **completion**.

1. Given a state-labeled rooted forest *ℱ* = (*F, ℓ*_*F*_), fix an initial active-root list 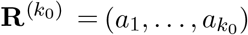 and initialize the output graph as *F*.
2. Compute 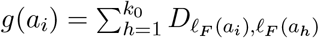 once for every initial component root.
3. While the active list contains *k >* 1 roots:
4. If *k* = 2, select the sole pair. Otherwise, compute 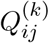 for every *i < j* as in Equation (1) and select a minimum-scoring pair using the seeded uniform choice among exact minimizers.
  a. Choose the member with smaller fixed score *g* as the parent root. Use an equiprobable seeded choice when the scores are equal.
  b. Add the directed parent-root–to–child-root edge with their dissimilarity as its weight.
  c. Delete the child root from the active list while preserving the relative order of all surviving roots.
5. Return the augmented tree and the sole surviving active root. If *k*_*0*_ = 1, return *F* and its component root unchanged.

#### Proposition 5.

*Algorithm S1 returns a closed-state top completion of ℱ without introducing a vertex. At every reduction, the active roots are the roots of pairwise vertex-disjoint components, and their dissimilarity matrix is the restriction of the initial component-root matrix to the surviving roots. For 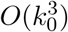 initial components, the algorithm uses 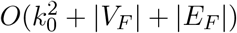arithmetic operations and memory*.

*Proof*. Initially the statement holds because the components of a forest are vertex-disjoint. A reduction adds an edge from the root of one component to the root of another. The parent remains a component root, the child acquires exactly one incoming edge, and no edge internal to either component is changed. The operation therefore joins two components without creating a directed cycle. Deleting only the active child root leaves every dissimilarity between surviving roots equal to its initial entry. Induction over the *k*_*0*_ − 1 reductions yields one rooted tree containing *F*. All added edges enter former component roots, and the labeling remains *ℓ*_*F*_. Hence the result is a closed-state top completion.

The fixed orientation scores require 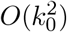 work. With *k* active roots, forming and searching the *Q* scores and restricting the active matrix require *O*(*k*^*2*^) work. Summing over *k* = *k*_*0*_, *k*_*0*_ − 1, …, 2 gives 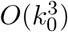 time. The initial and active matrices require 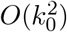 entries. The remaining storage is the forest and its *k*_*0*_ − 1 added edges.

### 9.9 Computational complexity

The following bounds use unit-cost arithmetic and count stored profile entries. Write *n* = |*U* | and *o* = |*O*|. For CTBF, *D* is supplied, so these bounds exclude its construction.

The pooled methods *M*_*0*_, *M*_*1*_, *M*_*3*_, and *M*_*5*_ require *O*(*n*^*3*^ + *mo*) time and *O*(*n*^*2*^ + *mo*) space, including input-profile storage and preparation. At *k* active roots, pair selection and matrix reduction take *O*(*k*^*2*^) work. Summing over the reductions gives *O*(*n*^*3*^). The profile-based orientation checks in *M*_*5*_ add *O*(*mn*) work. Storage for matrices and pair-index arrays remains quadratic throughout the reductions.

For *M*_*2*_, *M*_*4*_, *M*_*6*_, and 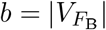 be the bottom-forest size after deferred copying and 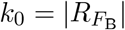 its number of roots. Conservative bounds are

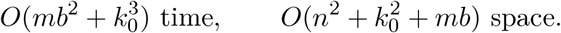

The bottom construction examines at most *O*(*b*^*2*^) candidate pairs, with *O*(*m*) work per plausibility check. Top completion takes 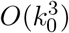 pair-selection work and adds at most *k*_*0*_ − 1 nodes. Each initialized occurrence can generate a chain of at most *q* − 1 deferred copies. Consequently,

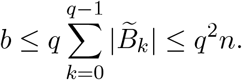

Copying therefore preserves a polynomial bound in the input dimensions. The partial projection of *M*_*2*_ adds only linear work.

MEDICC2 receives *o* occurrence profiles and a diploid outgroup. For the fixed total-copy-number, no-WGD configuration used here, let *T*_*MED*_(*m*) and *S*_*MED*_(*m*) bound the time and auxiliary space of one minimum-event distance calculation. Its serial topology-only pipeline then requires

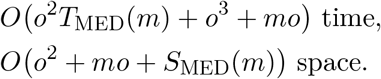

These bounds separate the quadratically many distance evaluations from the cubic NJ stage and do not assume a constant cost for event-distance calculation. Measured distance-inclusive pipeline resources are reported in Table 11.

### 9.10 Exact temporal-arborescence implementation

The temporal and no-time constructions are secondary occurrence-level methods and are not included in the principal synthetic comparison reported in the main Results. The temporal and no-time methods use, respectively, the admissible edge and root sets

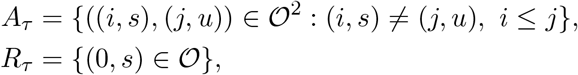

and

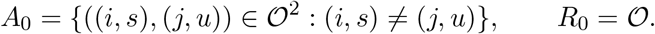

Define the occurrence labeling *l*_*O*_ : *O* → *U* by *l*_*O*_(*i, s*) = *s*. Thus the temporal method permits same-biopsy edges but excludes ancestry from a later to an earlier biopsy. For *ω* ∈ *{τ*, 0*}*, let *T* = (*O, E*_*T*_) range over directed spanning trees with *E*_*T*_ ⊆ *A*_*ω*_ and root *r* = (*i*_*r*_, *s*_*r*_) ∈ *R*_*ω*_. The method minimizes lexicographically

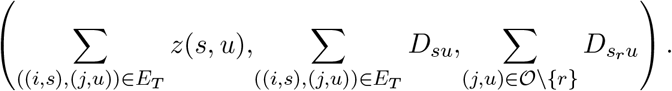

A seeded ranking of candidate edges and roots is the fourth coordinate among solutions tied on all three scientific criteria. Canonical order resolves any remaining equality. The temporal method uses *ω* = *τ*, whereas the no-time method uses *ω* = 0. Both return (*T, l*_*O*_), retain one labeled node for every member of *O*, and introduce no additional node. For the temporal method, the identity map on *O* is an observation map satisfying temporal compatibility. For the no-time method, it represents every observation but need not satisfy temporal compatibility.

Let *n* = |*O*|. For the implementation, occurrences are placed in a canonical order by lexicographic CNP, followed by biopsy index for repeated occurrences of the same state. This order assigns the occurrence indices 0, …, *n* − 1. The complete tie universe consists, in parent-major order, of all *n*(*n* − 1) directed pairs of distinct occurrences followed by the *n* possible roots. A permutation initialized by the reported seed assigns a distinct integer rank to every member of this *n*^*2*^-element universe. The temporal and no-time methods use the same complete universe, so every candidate shared by the two modes receives the same rank. Among solutions tied on the three scientific coordinates above, the fourth coordinate is the sum of the ranks *ρ*(*e*) of the selected biological edges and the rank *ρ*(*r*) of the selected root. A remaining equality of rank sums is resolved by the canonical candidate order.

#### 9.10.1 Exact integer scalarization and virtual root

Every stored floating-point dissimilarity is converted to its exact rational value and represented as an integer under one common denominator. Let *δ*_*su*_ denote the resulting integer unit for *D*_*su*_, let *δ*_*max*_ = max_*s,u*∈*U*_ *δ*_*su*_, and let *ρ*_*max*_ = *n*^*2*^ − 1 be the greatest seeded rank. Write a candidate root as *r* = (*i*_*r*_, *s*_*r*_). For an augmented representation containing exactly one virtual-root edge, upper bounds on the four lower-coordinate totals are

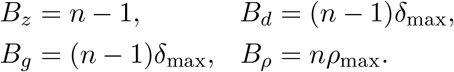

Define the mixed-radix coefficients

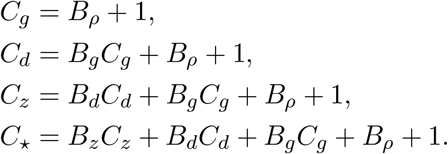

For an admissible biological edge *e* = ((*i, s*), (*j, u*)) ∈ *A*_*ω*_, assign the integer cost

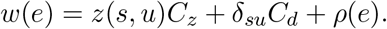

Introduce a virtual root *v*_⋆_. For every admissible biological root *r* ∈ *R*_*ω*_, add the edge *v*_⋆_ → *r* with cost

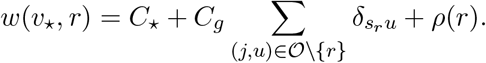

The augmented graph is rooted at *v*_⋆_. Any spanning arborescence of this graph contains at least one virtual-root edge. A feasible biological arborescence augmented by one such edge has total cost strictly less than 2*C*_⋆_, whereas every augmented arborescence containing at least two virtual-root edges has cost at least 2*C*_⋆_. Hence a minimum-cost arborescence contains exactly one virtual-root edge whenever a feasible biological arborescence exists. Its target is the selected biological root. Removing *v*_⋆_ and this edge returns a feasible arborescence on *O*.

##### Proposition 6.

*Minimizing the scalar cost of the augmented arborescence is equivalent to lexicographically minimizing the irreversible-zero violations, total edge dissimilarity, root score, and seeded rank sum, in this order*.

*Proof*. After the virtual-root argument above, every optimum contains exactly one virtual-root edge. Its four lower-coordinate totals are nonnegative integers bounded by *B*_*z*_, *B*_*d*_, *B*_*g*_, and *B*_*ρ*_, respectively. By construction, one unit multiplied by *C*_*g*_ exceeds every possible rank sum. One unit multiplied by *C*_*d*_ exceeds every possible root-score and rank contribution. One unit multiplied by *C*_*z*_ exceeds every possible lower-coordinate contribution. Comparison of two feasible scalar costs consequently agrees with lexicographic comparison at the first coordinate on which the corresponding arborescences differ. □

#### 9.10.2 Cycle-contraction solver and complexity

The augmented integer-weighted graph is solved by a compact Chu–Liu/Edmonds minimum-arborescence algorithm [3, 8]. For every nonroot vertex, the solver first selects a minimum-weight incoming edge, taking the first edge in canonical iteration order when integer weights are equal. If the selected edges contain no directed cycle, they form the required arborescence. Otherwise, one selected directed cycle *C* is contracted. Let *e*_*v*_ be the selected edge entering a cycle vertex *v* and let *M* be the maximum of *w*(*e*_*v*_) over the vertices *v* of *C*. An edge of weight *w*(*x, v*) entering the cycle is assigned the contracted weight

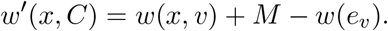

Edges leaving the cycle retain their weights, and edges internal to the cycle are omitted from the contracted graph. The procedure continues on the contracted graph. During reverse expansion, the selected edge entering a contracted component identifies one cycle vertex. Every selected cycle edge except the edge entering that vertex is restored. The cycle-contraction optimality argument of Chu–Liu and Edmonds then yields a minimum-cost spanning arborescence of the augmented graph.

The occurrence graph contains *O*(*n*^*2*^) candidate biological edges and at most *n* virtual-root edges. Constructing their plausibility costs requires *O*(*mn*^*2*^) profile-entry operations. For the conventional cycle-contraction procedure, the number of graph operations is proportional to the number of vertices times the number of edges and is therefore *O*(*n*^*3*^). The implementation stores one mutable edge universe, the occurrence-level matrix data, and linear-size contraction records, giving *O*(*n*^*2*^) stored entries in addition to the CNP profiles. These bounds count arithmetic and stored objects. The bit complexity of the exact integer operations also depends on the binary representations of the input dissimilarities.

## 10 Supplementary experimental methods

### 10.1 Simulation and biopsy collection

Table 3 specifies the standard regime.

The initiation scale is a phenomenological approximation to ongoing or post-burst diversification in patient-derived colorectal-cancer organoids, for which Bollen et al. observed approximately one CNA event per ten cell divisions [2]. The correspondence is phenomenological: simulation counts initiation proposals on attempted lineage transitions, whereas the study counts cell-division events. The crucial mask contains the zero-based simulator indices *{*4, 7, 38, 40, 41, 49, 54, 61, 69, 72*}*. Its 10% size is a coarse viability prior motivated at the order-of-magnitude level by cancer-cell-line dependency screens [5]. The synthetic bins have no gene-level annotation.

For each successful replicate, one history is generated to height 34. At each *H* ∈ *{*14, 24, 34*}*, its topology and profile assignment are restricted to *T* ^⋆^|_≤*H*_. The restricted history is regarded as a CNP-tree over the image of that profile assignment, with state labels and the profile-to-label bijection inherited by restriction. This CNP-tree is the paired true tree at depth *H*. The three-biopsy placement policies are defined by

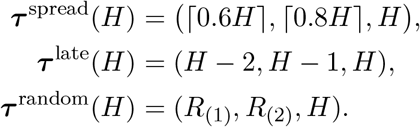

For the random policy, *R*_*1*_ and *R*_*2*_ are sampled uniformly without replacement from the integer set

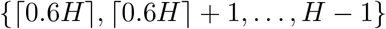

using a dedicated placement seed, and *R*_(1)_ *< R*_(2)_ are their order statistics. Thus the spread schedule places observations at nominal fractions 60%, 80%, and 100% of the history, the late schedule uses the final three generations, and the random schedule varies the first two observations within the same late-time window while fixing the terminal biopsy.

The standard three-biopsy schedule is ***τ*** ^*spread*^. It gives schedules (9, 12, 14), (15, 20, 24), and (21, 28, 34) for the three respective heights. Five prespecified sensitivity conditions are derived from the height-24 prefix. Biopsy-number schedules are

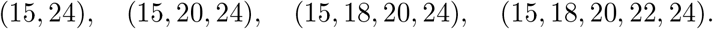

The placement comparison holds the number of biopsies at three and compares the standard schedule (15, 20, 24) with the late schedule (22, 23, 24) and a random schedule (*R*_(1)_, *R*_(2)_, 24). Here *R*_*1*_ and *R*_*2*_ are drawn uniformly without replacement from *{*15, …, 23*}* using a dedicated placement seed, and *R*_(1)_ *< R*_(2)_ are their order statistics. Thus every history provides eight prespecified experimental conditions. A random schedule may coincide with another schedule.

If selected generation *τ*_*k*_ contains *N*_*k*_ *>* 0 retained genotype-state occurrences, its sample size

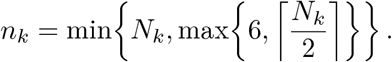

The *n*_*k*_ occurrences are sampled uniformly without replacement using the prespecified biopsy seed. Conditions sampling the same generation within one history use the same sampled occurrence set. Their distinct state labels form *B*_*k*_, and their true-tree nodes define the sampling map *σ*. An empty scheduled generation is a failed replicate and is not replaced. Because sampling is over retained genotype-state occurrences, the 50% fraction is an observation-density parameter rather than a tissue or clone fraction.

### 10.2 Dissimilarity construction

Let 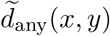 denote the directed event count returned by the improved `cnp2cnp` approximation in any mode for source profile *x* and target profile *y* [4]. For every observed state pair *s, t* ∈ *U*, we compute the two directed `cnp2cnp` costs and use the symmetric dissimilarity

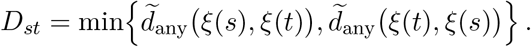

The underlying event model permits one segmental amplification or deletion to change copy number by more than one [4]. Taking the minimum symmetrizes the potentially unequal directed costs. This minimum-of-directions construction is the one reported for the original cnp2cnpNJ experiments [4]. The released matrix-mode routine^1^ instead evaluates only 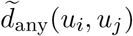 for *i < j* and mirrors that value. Its matrix can therefore change with FASTA input order when the two directed costs differ, and the NJ topology can consequently change. Evaluating both directions makes CTBF’s matrix invariant to profile order. For each condition, *D* is defined on its observed state set *U*. The frozen confirmation run computed one superset matrix on the union of the eight state sets within each history block and supplied each condition with its exact label-aligned submatrix. This batching changes neither an entry of *D* nor an inferred tree because every entry depends only on its profile pair. A separate audit reconstructed the 300 standard matrices directly from their condition-specific state sets and reproduced every frozen matrix and *M*_*2*_ labeled topology exactly. A failed distance calculation is recorded as an input failure, and no alternative dissimilarity is substituted.

### 10.3 Evaluation metrics

Before computing a score, we verify that the true and reconstructed graphs are nonempty rooted trees with exactly one root, parent-to-child edge orientation, and at least one labeled node. Every reconstructed state label must belong to *U*. Observation-label coverage and all validation failures are recorded explicitly. Complete coverage is required for analysis, so an invalid or incomplete reconstruction is retained as a failed record.

#### 10.3.1 Ancestor–descendant F1

For a CNP-tree or partial CNP-tree *T* = (*T, l*), define the set of unique ordered state-label relations

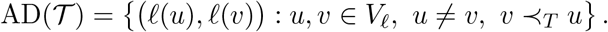

The first element labels the ancestor. Relations induced by different occurrence pairs but having the same ordered labels contribute once, while (*s, s*) is retained when one occurrence of state *s* is ancestral to a distinct occurrence of the same state. Define

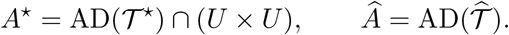

The corresponding precision and recall are

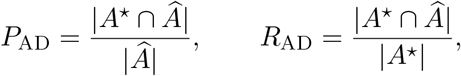

with value zero whenever the respective denominator is zero. We report

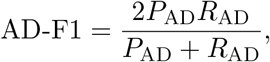

also with value zero when its denominator is zero. AD-F1 measures directed state-label ancestry, not direct-edge recovery or correspondence between repeated occurrences. Unlabeled internal nodes contribute no labels to this relation set. AD-F1 is therefore primary for closed-state reconstruction and diagnostic only for partial reconstruction.

#### 10.3.2 Rooted generalized Robinson–Foulds similarity

We use the generalized Robinson–Foulds construction for multi-labeled evolutionary trees [18]. Every node *v* ∈ *V*_*T*_ induces the multiset

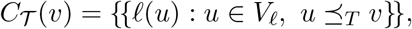

and the tree representation *C*(*T*) = *{{C*_T_ (*v*) : *v* ∈ *V*_*T*_ *}}* retains both label multiplicities within clusters and the multiplicity of equal clusters. For cluster multisets *a* and *b*, let *a*_*s*_ and *b*_*s*_ denote the multiplicities of label *s* in *a* and *b*, respectively, and let

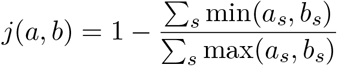

be their multiset-Jaccard distance. Every leaf is labeled in a valid partial CNP-tree or CNP-tree. Every node is a leaf or has a descendant leaf. Consequently, each cluster contains at least one label, and the denominator in *j* is positive. Write 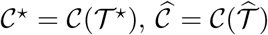, and 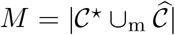, where multiset union and difference retain multiplicity. The implemented rooted GRF distance is

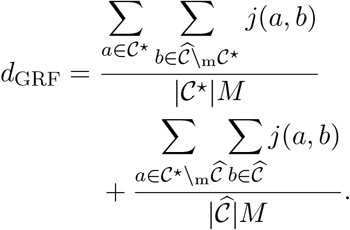

We report the similarity GRF = 1 − *d*_*GRF*_, for which larger values indicate greater agreement. Rooted GRF is primary for partial reconstruction and complementary for closed-state reconstruction. It compares the complete true and reconstructed cluster multisets, so unobserved true states, tree-size differences, and repeated labels all contribute to the score. For *M*_*0*_, rooted GRF is computed on the returned synthetic-root representation. This permits evaluation under the common rooted partial-tree endpoint, but it is neither a root-recovery assessment nor a reproduction of the unrooted Robinson–Foulds comparison in the cnp2cnp study.

#### 10.3.3 State-restricted true-tree evaluation

##### Definition 7

(Trimmed true tree). For the observed state set *U*, let

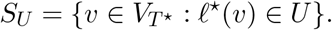

The *trimmed true tree* 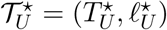is obtained from *T* ^⋆^ by removing the labels of all nodes outside *S*_*U*_ and applying the following reductions until none is applicable:

1. Delete an unlabeled leaf and its incident edge.
2. Suppress an unlabeled nonroot node with exactly one child by joining its parent directly to its child.
3. If the root is unlabeled and has exactly one child, delete the root and promote its child to the root.

Every node in *S*_*U*_ is retained, with 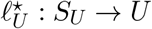 given by 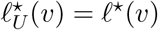.

Equivalently, 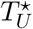 is the minimal connected subtree of *T* ^⋆^ spanning *S*_*U*_, rooted at its node of minimum original depth, with all unlabeled nodes having exactly one child suppressed. This characterization makes the result independent of the order of reductions. Since *U* is nonempty and observed in the true tree, the result is nonempty and every leaf is labeled. Every retained unlabeled node has at least two children. An edge of the trimmed tree can represent a path of the true tree rather than a single evolutionary transition.

The set *S*_*U*_ includes all true occurrences of observed states, not only the sampled occurrences *σ*(*O*). Labeled nodes with one child are retained, and distinct nodes with the same label are never merged. Consequently, unsampled occurrences of observed states, including same-state ancestral paths, still contribute to the evaluation. Figure 5 illustrates both distinctions. Trimming preserves ancestry among retained labeled nodes. Thus

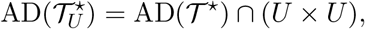

so the true relation set used by AD-F1 is unchanged.

Writing GRF(*T*_*1*_, *T*_*2*_) for the rooted similarity defined above, the two evaluation targets give

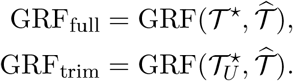

The reconstructed tree is identical in both comparisons. The first retains the complete true-history representation. The second restricts the truth to observed states while preserving their occurrence multiplicities and connecting topology. Neither score is an occurrence-matching accuracy measure. The partial-tree comparison reports both scores on the same reconstructed trees. The original primary test retains GRF_*full*_. Across the completed reevaluation, the observed-state AD relation sets and all AD-F1 fields agree exactly between targets.

### 10.4 Paired comparison protocol

Each prospectively seeded evolutionary history is one experimental block. All eight conditions derived from that history remain in the same block. For each condition, all eight CTBF methods receive the same biopsy sample and aligned dissimilarity matrix and use the same prespecified seed within that condition. Truth is unavailable during reconstruction and is loaded only by the evaluator. Every contrast is paired within blocks and remains within either the partial or closed-state output class.

In the paired comparisons below, a difference denoted by two method names means the first method’s score minus the second method’s score on the same input. The sole prespecified primary comparison is *M*_*2*_ − *M*_*0*_ in rooted GRF. For each block, its effect is the unweighted mean of the paired score differences at heights 14, 24, and 34. A block enters this analysis only if both methods succeed at all three heights. The directional claim requires at least 90 complete blocks, an arm-specific failure rate no greater than 10% at every height, a mean block effect of at least 0.01, a positive lower endpoint of the two-sided 95% block-bootstrap percentile interval, and *p <* 0.05 in a one-sided block sign-flip test for a positive mean. Both procedures use 100,000 prespecified random draws. The sign-flip *p*-value uses the plus-one correction. Its null calibration assumes sign symmetry of the independent block-effect distribution. This is one primary test and therefore requires no multiplicity adjustment.

Prespecified secondary analyses comprise the height-specific absolute scores, the block-averaged AD-F1 contrasts *M*_*6*_ − *M*_*5*_ and *M*_*7*_ − *M*_*3*_, the height-24 biopsy-number and placement comparisons, and a separate MEDICC2 comparison on the 300 standard conditions. Secondary intervals use history-block resampling and do not define additional binary conclusions. For a statistic based on *n* eligible independent histories, each bootstrap replicate samples *n* histories with replacement and recomputes that statistic from their stored scores. All method and condition scores of a selected history remain together. For a paired contrast, the statistic is computed from within-history differences. Combined-height contrasts first average those differences within each history. The 2.5th and 97.5th percentiles of 100,000 bootstrap replicates define the reported interval. These are pointwise intervals, rather than simultaneous confidence bands over all methods and conditions. They describe sampling uncertainty across histories under the specified simulation and sampling design. In complete-case comparisons, they are conditional on the eligible histories. Bootstrap replication neither generates new histories nor repeats biopsy sampling or reconstruction. The prespecified biopsy-number analysis used *M*_*6*_ to assess recovery as additional longitudinal biopsies were supplied. It reported the two-to-five-biopsy curve and the five-minus-two contrast, without a confirmatory decision rule or a requirement for adjacent monotonicity. Let *U* ^*(2)*^ be the observed state set in the two-biopsy condition of a given block. For every biopsy count, the fixed-label endpoint replaces *A*^⋆^ and *Â* by

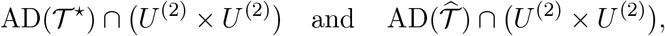

respectively, before computing AD-F1. All reconstructed nodes remain available when determining ancestry, but only relations with both labels in *U* ^*(2)*^ are scored.

The main biopsy-number figure uses ordinary AD-F1 for all five closed-state methods, scoring the observed-state set of each condition. The fixed-label endpoint is retained as a supplementary control, including the original *M*_*6*_ endpoint interval and the paired five-minus-two gain contrasts *M*_*6*_ − *M*_*5*_ and *M*_*7*_ − *M*_*3*_. The ordinary-score curves and the extensions to other methods are descriptive and introduce no additional hypothesis tests. Partial-tree GRF retains the complete reconstructed representation. Its trimmed-truth counterpart uses the true tree restricted to the observed state set of each condition. A further post hoc descriptive extension summarizes closed-state rooted GRF at the three standard heights and the corresponding *M*_*6*_ − *M*_*5*_ and *M*_*7*_ − *M*_*3*_ contrasts. It uses the stored scores for all five closed-state methods, with 100,000 paired history-block bootstrap draws and no additional hypothesis tests. Rooted GRF retains complete tree representations rather than the fixed-label ancestry target. The placement analysis prospectively assessed sensitivity to standard, late, and random schedules. It did not specify a directional hypothesis and is not interpreted as a causal effect under a fixed observed-state set. The simulated history is the independent unit. Edges, state pairs, biopsies, conditions, and reconstruction records are not treated as independent observations. We report attempted and complete denominators, absolute scores, paired mean and median differences, wins/ties/losses at tolerance 10^−12^, coverage, runtime, peak resident memory, and typed failures. Missing values are not imputed. Invalid outputs, incomplete label coverage, failed distance calculations, and resource-guard failures remain in the attempted count. No history or seed is replaced.

For the pipeline resource comparison, a separate audit processed each of the 300 standard conditions independently. It constructed the condition-specific matrix *D* and then ran *M*_*2*_. The *M*_*2*_ pipeline time is the sum of those sequential stages, and its peak resident memory is the larger stage peak. Exact matrix and labeled-topology equality were verified against the frozen confirmation output before resource values were summarized. MEDICC2 time and memory include its internally computed distances and topology. The *M*_*2*_ audit and MEDICC2 run both used four concurrent case workers. MEDICC2 itself used one core per case. Evaluation time and memory are excluded from both pipelines. The two totals therefore describe the executed pipelines without isolating either topology-construction complexity or biopsy information.

## 11 Supplementary results

Square brackets in the text denote pointwise 95% bootstrap intervals as defined in Supplementary Methods 10.4.

Before generating any confirmation history, we fixed the design, source inventory, simulator configuration, method roster, seeds, endpoints, and decision rule and recorded their checksums. The 100 new histories were disjoint from all method-development histories. The initial seven-method campaign completed all 800 input conditions and 5,600 CTBF reconstructions. Including *M*_*4*_ gives 6,400 successful CTBF reconstructions. All 300 MEDICC2 reconstructions also succeeded. No history, condition or method record was replaced. MEDICC2 was restricted to the three standard conditions from each history, whereas all eight CTBF methods were evaluated on all eight conditions.

### 11.1 Partial reconstruction and prespecified comparison

Table 4 gives absolute scores and paired effects. Figure 6 shows their primary history-block summary.

**Figure 6.**
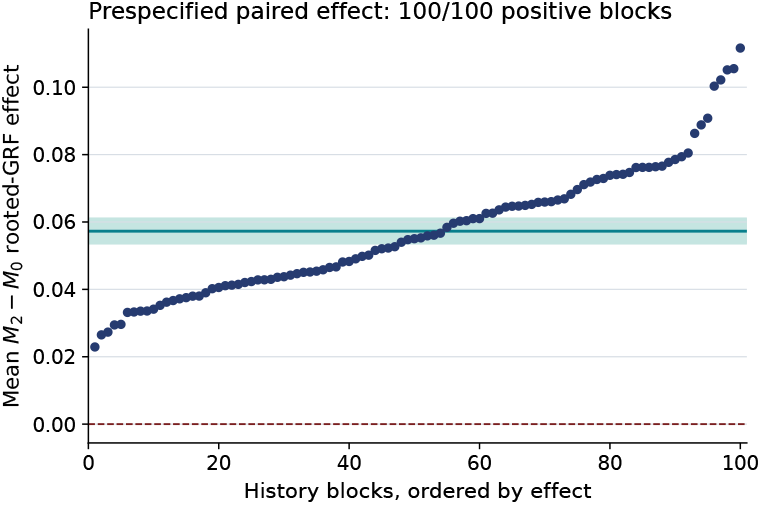
The 100 independent history-block effects for the prespecified *M*_2_ −*M*_0_ rooted-GRF comparison, ordered by effect. Each effect averages the three standard height-specific differences from one history. The horizontal line and shaded band show the mean and its two-sided 95% block-bootstrap interval. All block effects are positive.

For the prespecified *M*_*2*_ − *M*_*0*_ contrast, all 100 blocks were complete and the maximum primary-arm failure rate was zero. The median block effect was 0.055159. The one-sided 100,000-draw sign-flip test for a positive mean used the plus-one Monte Carlo correction and gave *p* = 9.9999 *×* 10^−6^, the smallest attainable value with this number of draws. All five prespecified support criteria were met. The condition-level wins/ties/losses were 298/0/2. MEDICC2 was a prespecified secondary comparator. Its condition-level comparison with *M*_*2*_ gave 275/0/25 wins/ties/losses in favor of *M*_*2*_.

### 11.2 Closed-state reconstruction

Table 5 reports absolute AD-F1 scores and the prespecified descriptive paired contrasts at each depth.

The paired AD-F1 effects change sign with depth, giving a depth-dependent ranking under this endpoint. These are the prespecified descriptive comparisons defined in Supplementary Methods 10.4.

Complementary rooted GRF gives a stable ordering of mean scores: *M*_*6*_, *M*_*7*_, *M*_*5*_, *M*_*1*_, and *M*_*3*_, from highest to lowest at each height (Table 6). Mean similarity decreases with depth for all five methods. The *M*_*6*_ −*M*_*5*_ mean effect decreases from 0.075185 at height 14 to 0.025705 at height 34. The corresponding *M*_*7*_ − *M*_*3*_ effects are 0.130706 and 0.067191. Both contrasts have positive three-height mean effects in every history block. These descriptive results concern complete cluster representations, including unobserved true states, repeated labels, and differences in tree size. AD-F1 remains the primary closed-state endpoint and measures ancestry among observed states. The different rankings reflect these distinct evaluation targets.

### 11.3 Biopsy-number and placement sensitivity

The biopsy-number comparisons use the same 100 height-24 histories. The two-, three-, four-, and five-biopsy schedules sample generations (15, 24), (15, 20, 24), (15, 18, 20, 24), and (15, 18, 20, 22, 24), respectively. Table 7 gives ordinary closed-state AD-F1 over the observed states of each condition. Tables 8 and 9 retain the fixed-two-biopsy-label control and the partial-tree full-truth GRF scores. Figure 7 shows this control and the trimmed-truth GRF curves for partial and closed-state reconstruction.

**Table 8.** Mean reconstruction scores by biopsy number at height 24. Each entry averages 100 histories. The closed-state AD-F1 target is fixed within each history, whereas partial-tree GRF includes changing observed-label coverage and tree representation.

| Method | 2 biopsies | 3 biopsies | 4 biopsies | 5 biopsies |
| --- | --- | --- | --- | --- |
| *Partial: full-truth rooted GRF* |  |  |  |  |
| $M_0$ | 0.100102 | 0.111363 | 0.114218 | 0.123468 |
| $M_2$ | 0.120594 | 0.154148 | 0.170419 | 0.210822 |
| $M_4$ | 0.119877 | 0.153401 | 0.169691 | 0.210320 |
| *Closed-state: fixed-label AD-F1* |  |  |  |  |
| $M_1$ | 0.490590 | 0.446632 | 0.446974 | 0.445120 |
| $M_3$ | 0.595484 | 0.602322 | 0.602939 | 0.609938 |
| $M_5$ | 0.746287 | 0.772461 | 0.784281 | 0.793432 |
| $M_6$ | 0.733913 | 0.769477 | 0.791224 | 0.826456 |
| $M_7$ | 0.663751 | 0.712812 | 0.734327 | 0.765809 |

**Table 9.**
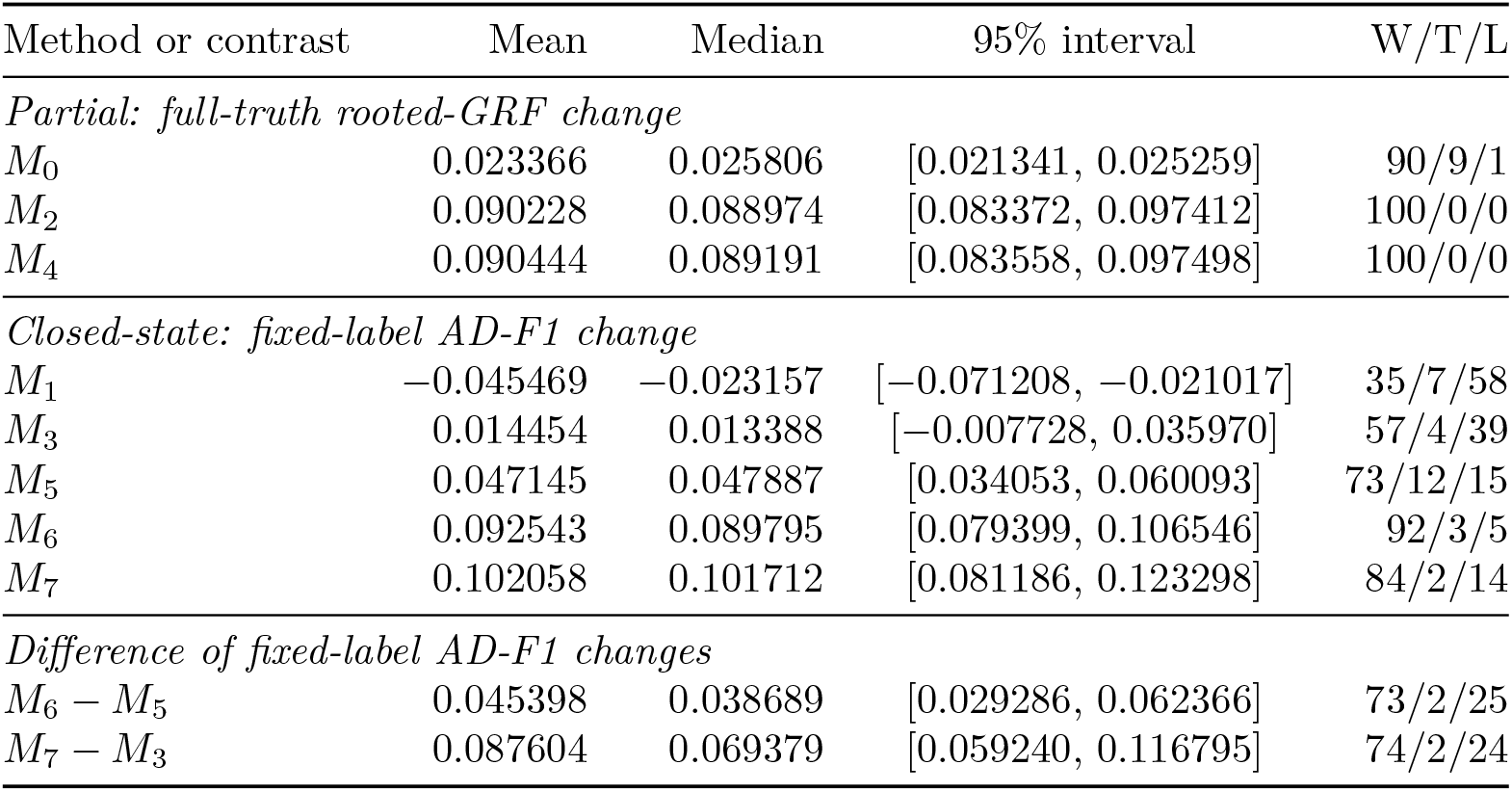
Paired five-minus-two-biopsy changes and differences between those changes. Each row contains 100 history-block effects. Positive values favor five biopsies, or the first method in a difference-of-changes row. All entries are descriptive.

| Method or contrast | Mean | Median | 95% interval | W/T/L |
| --- | --- | --- | --- | --- |
| *Partial: full-truth rooted-GRF change* |  |  |  |  |
| $M_0$ | 0.023366 | 0.025806 | [0.021341, 0.025259] | 90/9/1 |
| $M_2$ | 0.090228 | 0.088974 | [0.083372, 0.097412] | 100/0/0 |
| $M_4$ | 0.090444 | 0.089191 | [0.083558, 0.097498] | 100/0/0 |
| *Closed-state: fixed-label AD-F1 change* |  |  |  |  |
| $M_1$ | -0.045469 | -0.023157 | [-0.071208, -0.021017] | 35/7/58 |
| $M_3$ | 0.014454 | 0.013388 | [-0.007728, 0.035970] | 57/4/39 |
| $M_5$ | 0.047145 | 0.047887 | [0.034053, 0.060093] | 73/12/15 |
| $M_6$ | 0.092543 | 0.089795 | [0.079399, 0.106546] | 92/3/5 |
| $M_7$ | 0.102058 | 0.101712 | [0.081186, 0.123298] | 84/2/14 |
| *Difference of fixed-label AD-F1 changes* |  |  |  |  |
| $M_6 - M_5$ | 0.045398 | 0.038689 | [0.029286, 0.062366] | 73/2/25 |
| $M_7 - M_3$ | 0.087604 | 0.069379 | [0.059240, 0.116795] | 74/2/24 |

**Table 10.** Height-24 placement sensitivity. Entries are standard-schedule-minus-comparator mean effects, 95% block-bootstrap intervals, and wins/ties/losses. GRF is used for partial methods and AD-F1 for closed-state methods.

| Method | Metric | Standard minus late |  | Standard minus random |  |
| --- | --- | --- | --- | --- | --- |
| $M_0$ | GRF | -0.01564 | [-0.01751, -0.01375], | -0.00271 | [-0.00432, -0.00116], |
|  |  | 6/8/86 |  | 35/13/52 |  |
| $M_2$ | GRF | -0.04814 | [-0.05407, -0.04244], | -0.00809 | [-0.01307, -0.00325], |
|  |  | 3/2/95 |  | 37/2/61 |  |
| $M_4$ | GRF | -0.04840 | [-0.05446, -0.04263], | -0.00819 | [-0.01313, -0.00335], |
|  |  | 3/2/95 |  | 35/4/61 |  |
| $M_1$ | AD-F1 | -0.03639 | [-0.05761, -0.01499], | -0.00923 | [-0.03216, 0.01317], |
|  |  | 35/4/61 |  | 48/6/46 |  |
| $M_3$ | AD-F1 | 0.01576 | [-0.00591, 0.03759], | 0.00941 | [-0.00975, 0.02843], |
|  |  | 58/1/41 |  | 50/4/46 |  |
| $M_5$ | AD-F1 | 0.01538 | [-0.00479, 0.03576], | 0.01361 | [-0.00223, 0.02965], |
|  |  | 51/6/43 |  | 51/13/36 |  |
| $M_6$ | AD-F1 | -0.00883 | [-0.03222, 0.01525], | 0.00601 | [-0.01519, 0.02754], |
|  |  | 36/4/60 |  | 46/6/48 |  |
| $M_7$ | AD-F1 | -0.00107 | [-0.03097, 0.02780], | -0.00939 | [-0.03408, 0.01482], |
|  |  | 46/1/53 |  | 46/2/52 |  |

**Table 11.** Stored wall time and sampled peak resident memory. MB denotes 10^6^ bytes. The *M*_2_ pipeline includes condition-specific construction of its input matrix *D*.

| Stage | $n$ | Mean (s) | Median (s) | Max (s) | Mean MB | Max MB |
| --- | --- | --- | --- | --- | --- | --- |
| Condition-specific $D$ | 300 | 1.972 | 0.190 | 58.737 | 89.1 | 166.3 |
| $M_2$ reconstruction | 300 | 0.056 | 0.025 | 0.892 | 64.6 | 172.0 |
| $M_2$ pipeline | 300 | 2.028 | 0.216 | 59.629 | 89.1 | 172.0 |
| MEDICC2 pipeline | 300 | 10.332 | 6.612 | 139.925 | 268.0 | 453.0 |

**Figure 7.**
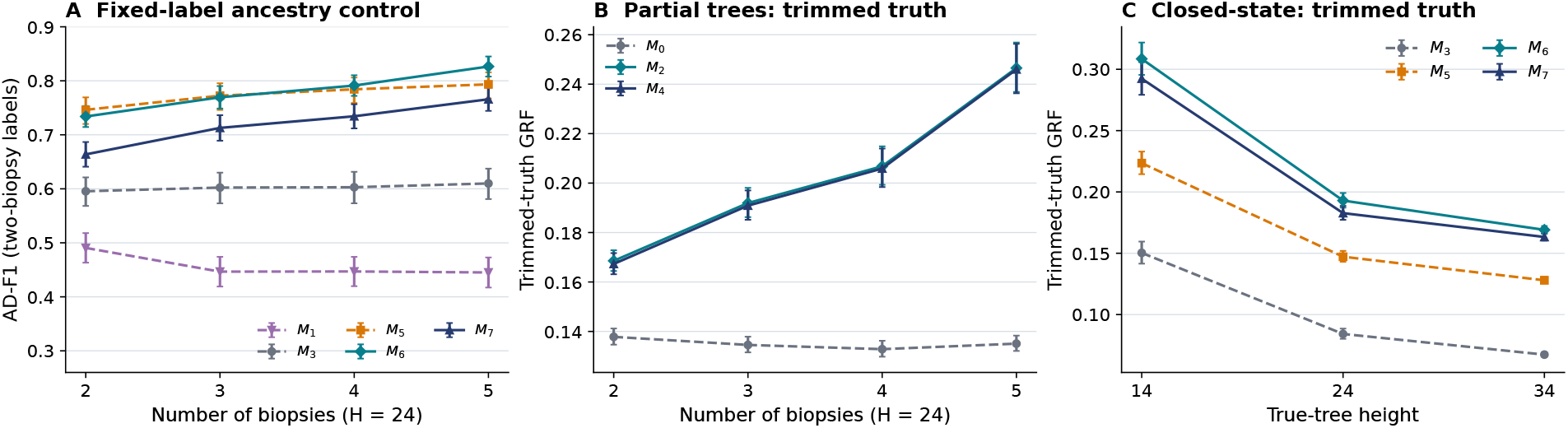
Sampling and truth-target controls, each based on all 100 histories. **(A)** Closed-state AD-F1 restricted in both trees to labels observed with two biopsies. Reconstruction uses all observations available in the corresponding condition. **(B)** Partial-tree trimmed-truth GRF by biopsy number at height 24. The trimmed true tree is determined separately by each condition’s observed-state set. **(C)** Closed-state trimmed-truth GRF by height. Solid and dashed curves denote biopsy-guided and pooled methods. Error bars are pointwise 95% whole-history bootstrap percentile intervals. AD-F1 is invariant to trimming the true tree and therefore requires no separate trimmed-truth curve.

Both biopsy-guided closed-state methods gained more from additional biopsies than their pooled counterparts using the same completion rule (Table 9). The mean excess gain was 0.045398 for *M*_*6*_ over *M*_*5*_ and 0.087604 for *M*_*7*_ over *M*_*3*_. The gain for pooled *M*_*5*_ also demonstrates a benefit from supplying additional profiles without their collection order. These post hoc contrasts assess how each complete construction responds to additional biopsies while holding the scored true relation set fixed. They include the effect of additional profiles as well as each method’s use of collection order.

The placement analysis addressed a different prespecified descriptive question: how sensitive were the methods to replacing the standard schedule with late or random sampling times? This descriptive comparison allows both the observed state set and its ancestral relations to change with the sampling schedule. Table 10 reports the placement contrasts.

The placement effects differed across methods and metrics.

### 11.4 Pipeline resources

Table 11 separates distance construction, reconstruction, and the combined pipeline measurements.

The first two rows report the sequential stages of the direct condition-specific *M*_*2*_ audit. Pipeline time is their per-condition sum, and pipeline memory is their maximum per-condition stage peak. MEDICC2 is timed as one executable invocation per standard condition. CTBF module imports precede its measured stages, whereas MEDICC2 executable startup is included. The resource totals include distance construction and reconstruction through these respective interfaces, with the stated difference in startup accounting. All 300 audited matrices and *M*_*2*_ labeled topologies matched the frozen confirmation output exactly.

### 11.5 Extended partial-tree comparison

We compared *M*_*0*_, *M*_*2*_, *M*_*4*_, MEDICC2 and NestedBD-Long on the same 100 independently generated histories. Method *M*_*4*_ shares the adaptive/deferred bottom with *M*_*2*_ and uses classical NJ top completion. All 800 *M*_*4*_ reconstructions succeeded. The full/trimmed scores of *M*_*0*_, *M*_*2*_ and MEDICC2 were reused without alteration, and the additional methods were evaluated with the same metric definitions. Table 12 reports both truth targets. Every NestedBD-Long input contributes the mean of its two run scores only when both are successful. This gives 100 paired inputs at height 14 and 98 at height 24. Each displayed five-method mean uses this same set. The four-method height-34 comparison retains all 100 histories.

**Table 12.** Matched partial-tree scores against full and trimmed truth. Each row uses the same successful histories for all available methods. NestedBD-Long scores are the means of both independently seeded runs. Its two incomplete height-24 inputs are excluded from every method in the height-24 rows. Height 34 contains the four available methods. The complete 100-history full-truth scores for *M*_0_, *M*_2_, *M*_4_ and MEDICC2 are given in Table 4.

| Target | $H$ | $n$ | $M_0$ | $M_2$ | $M_4$ | MEDICC2 | NestedBD-Long |
| --- | --- | --- | --- | --- | --- | --- | --- |
| Full | 14 | 100 | 0.18808 | 0.28526 | 0.28455 | 0.24700 | 0.27256 |
| Full | 24 | 98 | 0.11165 | 0.15452 | 0.15376 | 0.13420 | 0.12668 |
| Full | 34 | 100 | 0.08904 | 0.12089 | 0.12049 | 0.10259 | — |
| Trimmed | 14 | 100 | 0.19820 | 0.30620 | 0.30537 | 0.26198 | 0.28993 |
| Trimmed | 24 | 98 | 0.13474 | 0.19221 | 0.19115 | 0.16336 | 0.15379 |
| Trimmed | 34 | 100 | 0.11896 | 0.16817 | 0.16760 | 0.13851 | — |

**Table 13.** Additional paired partial-tree comparisons. NBD denotes NestedBD-Long. Each entry gives the mean score difference and its 95% whole-history bootstrap interval. W/T/L counts complete histories. Multi-height rows first average differences within each history. The full-truth primary *M*_2_ −*M*_0_ result remains in Table 4.

| Contrast | $H$ | $n$ | Full<br>mean [95% CI] | W/T/L | Trimmed<br>mean [95% CI] | W/T/L |
| --- | --- | --- | --- | --- | --- | --- |
| $M_2 - M_4$ | 14/24/34 | 100 | 0.00062<br>[0.00047, 0.00076] | 91/0/9 | 0.00081<br>[0.00065, 0.00097] | 94/0/6 |
| $M_4 - M_0$ | 14/24/34 | 100 | 0.05665<br>[0.05296, 0.06046] | 100/0/0 | 0.07071<br>[0.06718, 0.07438] | 100/0/0 |
| $M_4 - \text{MEDICC2}$ | 14/24/34 | 100 | 0.02501<br>[0.02274, 0.02735] | 100/0/0 | 0.03343<br>[0.03081, 0.03608] | 100/0/0 |
| $M_2 - \text{NBD}$ | 14 | 100 | 0.01271<br>[0.00732, 0.01805] | 69/8/23 | 0.01627<br>[0.01017, 0.02237] | 70/8/22 |
| $M_2 - \text{NBD}$ | 24 | 98 | 0.02784<br>[0.02459, 0.03111] | 93/2/3 | 0.03843<br>[0.03413, 0.04275] | 93/2/3 |
| $M_2 - \text{NBD}$ | 14/24 | 98 | 0.02011<br>[0.01689, 0.02328] | 86/2/10 | 0.02702<br>[0.02312, 0.03088] | 88/2/8 |
| $M_4 - \text{NBD}$ | 14/24 | 98 | 0.01937<br>[0.01615, 0.02250] | 86/2/10 | 0.02608<br>[0.02221, 0.02986] | 87/2/9 |

Combined-height effects average differences within a history before resampling. Descriptive percentile intervals use 100,000 whole-history bootstrap draws. For each height-specific figure panel, the same resampling indices are applied to all methods and both truth targets. Neither MCMC seeds nor multiple heights are independent histories. The original primary full-truth comparison is unchanged. No new hypothesis test or automatic method-selection rule is introduced.

The two biopsy-guided partial methods have the highest mean similarity at each evaluated height under both truth targets. The small *M*_*2*_ − *M*_*4*_ differences contrast with their much larger gains over pooled NJ (Table 13). Across the three standard heights, the trimmed-truth *M*_*2*_ − *M*_*0*_ effect is 0.07153 [0.06800, 0.07517], and the corresponding *M*_*2*_-minus-MEDICC2 effect is 0.03425 [0.03165, 0.03689]. Condition-level W/T/L over all 300 standard inputs is 298/0/2 for *M*_*2*_ − *M*_*0*_ under either target. For *M*_*2*_ versus MEDICC2 it is 275/0/25 under full truth and 280/0/20 under trimmed truth. For *M*_*2*_ − *M*_*4*_, the corresponding counts are 232/49/19 and 227/49/24. These counts complement the history-level summaries and do not increase the number of independent histories.

At height 14, *M*_*2*_ versus NestedBD-Long gives full/trimmed W/T/L of 69/8/23 and 70/8/22. At height 24, both targets give 93/2/3. Thus all 198 available input pairs give 162/10/26 and 163/10/25. The combined-height interval in Table 13 instead uses the 98 histories complete at both heights, averaging their two differences before resampling. NestedBD-Long has higher mean scores than MEDICC2 at height 14, whereas MEDICC2 has higher mean scores at height 24.

Table 14 complements the full-truth biopsy-number analysis. The two biopsy-guided methods gain under both targets, while pooled NJ’s five-minus-two contrast changes sign. Because additional biopsies change *U* and hence the trimmed target, these contrasts describe the complete sampling-and-reconstruction setting.

**Table 14.**
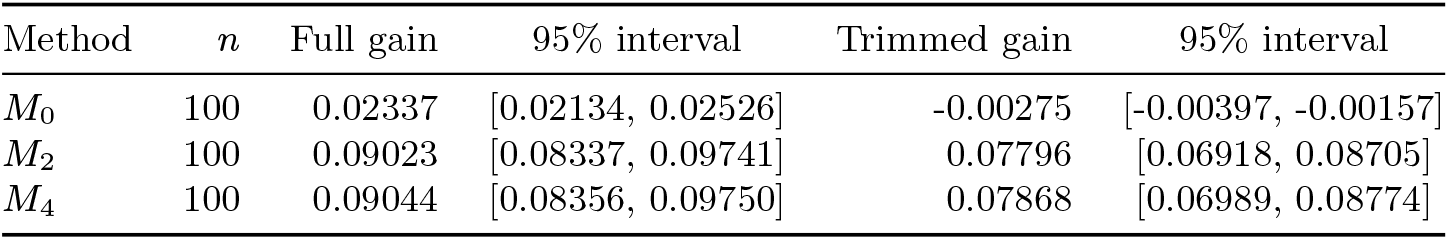
Five-minus-two-biopsy GRF differences at height 24. All 100 histories are paired. The trimmed true tree is constructed separately for each observed-state set, so these are not fixed-target sampling effects.

### 11.6 Preliminary synthetic NestedBD-Long analyses

Preliminary analyses used one development history, separate from the 100 confirmation histories below. An initial lossless pilot comprised two two-million-iteration runs at each of heights 14, 24, and 34, retaining all 100 bins and respectively 17, 60, and 288 biopsy-indexed occurrences. The fixed mean clock was initially one per simulator generation. All six runs completed, but the scalar and between-run tree diagnostics did not establish convergence. No reconstruction-accuracy comparison was made from these initial outputs.

A subsequent clock-only sensitivity analysis used the same height-14 and height-24 inputs, two seeds, and three fixed rates: 0.00026546875, 0.001061875, and 0.0042475 per generation. All twelve two-million-iteration runs completed. The middle rate was specified from the simulator’s proposal scale, as derived below, before the final comparison. The root exposure, demographic and sampling priors, heating, and input profiles were unchanged. At this nominal rate, posterior effective sample sizes after the fixed 10% burn-in were 81.3 and 98.9 at height 14, and 29.5 and 26.1 at height 24. Convergence remained unestablished.

A fixed-budget topology comparison then summarized both nominal-rate runs using the same sampled-ancestor-aware rule as the later comparison. At both depths, each external rooted-GRF score lay between the corresponding *M*_*0*_ and *M*_*2*_ scores. These paired depths provide preliminary evidence from one development history, separate from the confirmation histories below.

A separate budget diagnostic reran the first two confirmation histories at heights 14 and 24, with both seeds, for ten million iterations. Only the chain length changed. At the same 10% burn-in, posterior ESS reached 200 in seven of eight ten-million-iteration runs, compared with none of their eight earlier two-million-iteration counterparts. The first history at height 24 retained a maximum between-seed clade-frequency difference of 0.251. Longer runs improved these diagnostics without establishing convergence across all inputs. The 100-history comparison below uses a uniform two-million-iteration budget. The separate longer diagnostic is not mixed into its accuracy summaries.

### 11.7 Finite-budget NestedBD-Long comparison on synthetic inputs

**Design and inputs**. We added NestedBD-Long [17] as a contextual comparator after registration of the confirmation experiment. The completed comparison includes all 100 original histories. Each contributed the standard height-14 and height-24 conditions, sampled at generations (9, 12, 14) and (15, 20, 24), respectively. The existing 50% sampling rule with lower bound six was unchanged. All 200 inputs retained their original 100 bins and every biopsy-indexed occurrence. Height-14 inputs contained 3–38 occurrences and 1–25 distinct states. Height-24 inputs contained 3–190 occurrences and 1–150 distinct states. Copy numbers ranged from 0 to 6, within the tested external state range 0–9. No input was clipped, thinned, expanded or replaced. All-diploid inputs were retained. All 400 seed runs were executed for this uniform-budget comparison. The earlier ten-history results remain a separate preliminary analysis.

#### Inference and time scale

We used BEAST 2.6.7 with the unchanged released packages and the pinned NestedBD-Long source and XML cleanup described in Supplementary Methods 12.4. The cleanup removed the unused ucldMean state, its scale operator and log entry, and the joint parameter–tree up–down operator. It changed the sampled state space and proposal schedule, not the inference source or binary. Numerical sampling generations were supplied as times. The actual mean clock, meanClockRate, was fixed at 0.001061875 per generation before this comparison. This value matched the simulator’s first-order proposal quadratic variation per bin at copy number two to a critical linear birth–death reference process. Under the declared simulation parameters, the mean interval length is 25.75 bins and its signed copy-change second moment is 1.30. The resulting quadratic variation is 0.001[0.90 + 0.10(25.75)(1.30)] = 0.0042475 per bin and generation. Dividing by the reference coefficient 2 *×* 2 = 4 gives the fixed clock. This approximation matches the proposal-scale quadratic variation. Event overlap, cancellation, zero-copy absorption, viability filtering, selection, and the simulator’s nonzero drift remain outside this calibration. The separate diploid-root exposure OrigTime remained one and was not rescaled by this clock. The remaining origin, diversification, sampling, and heating settings were unchanged.

Each input received two independently seeded runs, each with four coupled chains and two million iterations, following a 10,000-iteration technical check. Each long run had a one-hour limit, a 2-GiB Java heap, and a 4-GiB process-tree resident-memory limit. At most two runs executed concurrently. Of 400 runs, 397 completed and produced valid scored tree summaries. At height 24, history 016 lost one run when BEAST’s posterior consistency check failed at iteration 9. The logged recomputation differed by approximately 0.01085, and the process exited with code 1. Both runs for history 085 exceeded the one-hour limit. These three failures were retained without replacement or numerical repair. Accuracy comparisons requiring both runs therefore include 100 height-14 and 98 height-24 inputs. The successful mate of history 016 remains in the diagnostics but does not supply a single-run accuracy substitute. The cold chain was sampled every 1,000 iterations. The fixed 10% burn-in discarded the first 200 of 2,001 samples, leaving 1,801 trees per run. The sampled-ancestor-aware BEAST `TreeAnnotator` selected a maximum-clade-credibility tree from each retained stream, keeping its sampled node heights. An exact score tie selected the first retained tree. The summary rule was fixed before accuracy scoring. Both runs were reported without pooling posterior samples or selecting a preferred seed.

#### Topology mapping and paired evaluation

Each observation retained its exact state and biopsy-indexed identity. A sampled ancestor represented by a zero-length terminal branch was mapped to its parent vertex, removing only that terminal representation. The inferred root, other edges, recurrent states, and all observations were preserved. Ambiguous mappings would have been rejected rather than repaired. Latent vertices received no inferred CNP labels. An observed root was retained if present, rather than imposing the CTBF partial-tree root convention. We evaluated these unchanged trees against both the complete true prefixes and their trimmed true trees using the multiplicity-aware rooted GRF of Supplementary Methods 10.3. The previously computed *M*_*0*_, *M*_*2*_ and MEDICC2 scores were retained unchanged. Method *M*_*4*_ was evaluated on the same inputs and targets.

The matched analysis in Supplementary Results 11.5 averages both successful seed scores within each input. Combined-height effects then average the height-14 and height-24 differences within each of the 98 complete histories. Wins, ties and losses use absolute tolerance 10^−12^. All additional intervals use 100,000 whole-history bootstrap resamples. Missing pairs remain in the attempted denominators, without score imputation.

#### Reconstruction results

The matched five-method means, paired effects and both truth targets are reported in Tables 12 and 13. NestedBD-Long has higher mean similarity than pooled NJ at both heights. Methods *M*_*2*_ and *M*_*4*_ have higher mean similarity than NestedBD-Long. Per-input and per-seed scores, including every failed outcome, accompany the numerical results.

#### Diagnostics and interpretation

Effective sample sizes (ESS) were computed from the retained cold-chain traces with BEAST `LogAnalyser`. Between-run clades are sets of descendant observation identifiers, excluding singleton and complete-taxon sets. Zero-length sampled-ancestor branches are retained for this diagnostic. A majority clade occurs in more than half of a run’s retained trees. Table 15 summarizes the fixed 10% burn-in and two additional diagnostic windows. At 10% burn-in, median posterior ESS is 85.4 at height 14 and 39.6 at height 24. Only 25/200 and 4/197 successful runs, respectively, reach posterior ESS 200. The median maximum between-run clade-frequency difference is 0.039 at height 14 and 0.317 at height 24. Identical majority-clade sets occur in 74/100 and 25/98 paired inputs. These scalar and tree-distribution diagnostics leave convergence unestablished under the declared budget. Agreement of summary scores alone does not establish posterior agreement.

**Table 15.** NestedBD-Long diagnostics for all successful runs in the 100-history comparison. ESS columns count individual cold-chain runs. Clade-frequency columns compare the two runs for each complete input (100 at height 14 and 98 at height 24). For each input, the largest absolute between-run clade-frequency difference is computed. The last column gives its median and maximum across inputs. The 25% and 50% windows are diagnostic only. All scored trees use 10%.

| $H$ | Burn-in (%) | Runs | Posterior ESS median [range] | Posterior ESS $\geq 200$ | Median likelihood ESS | Clade difference median [maximum] |
| --- | --- | --- | --- | --- | --- | --- |
| 14 | 10 | 200 | 85.4 [11.3, 506.2] | 25 | 760.8 | 0.039 [0.470] |
| 14 | 25 | 200 | 72.9 [13.2, 439.8] | 22 | 614.2 | 0.045 [0.599] |
| 14 | 50 | 200 | 54.7 [7.3, 386.9] | 14 | 411.4 | 0.050 [0.572] |
| 24 | 10 | 197 | 39.6 [4.1, 427.0] | 4 | 117.4 | 0.317 [0.926] |
| 24 | 25 | 197 | 36.1 [4.0, 345.1] | 3 | 114.9 | 0.319 [0.941] |
| 24 | 50 | 197 | 26.0 [4.5, 264.4] | 2 | 90.0 | 0.334 [0.949] |

**Table 16.** KTN152 reconstruction dimensions and computational resources.

| Stage or method | States | Vertices | Unlabeled | Time (s) | Peak RSS (MiB) |
| --- | --- | --- | --- | --- | --- |
| Chromosome-aware distance | 76 | – | – | 4.197 | 884.9 |
| $M_0$ | 76 | 151 | 75 | 1.693 | 208.4 |
| $M_2$ | 76 | 148 | 40 | 1.725 | 226.8 |

The accuracy comparison characterizes the point estimates produced by this fixed-budget procedure. The external model differs from the simulator’s correlated interval changes, multigains and representative-genotype sampling. The clock scale is approximately matched, while demographic, sampling and root-exposure assumptions remain model-specific. NestedBD-Long uses numerical sampling times, whereas CTBF uses biopsy order. The successful external JVM runs took 0.42–47.62 minutes each. These measurements exclude technical checks, controller work and postprocessing. The matched condition-specific pipeline resource comparison in Table 11 concerns *M*_*2*_ and MEDICC2.

## 12 Supplementary KTN152 empirical application

### 12.1 Public data and copy-number processing

The KTN152 data correspond to patient P14 in the longitudinal triple-negative breast cancer cohort of Kim et al. [16]. We used the single-nucleus DNA-sequencing libraries from pretreatment, after two cycles of neoadjuvant chemotherapy, and from surgery after six cycles. Raw reads are publicly available through BioProject PRJNA396019, also indexed as SRA study SRP114962. The archive contained 135 libraries annotated as individual KTN152 cells and three separately identified population libraries. We selected the individual-cell libraries for single-cell reconstruction. A prespecified minimum of 300,000 raw reads retained 132 cells for initial processing. A stricter technical threshold of one million reads retained 129 cells, distributed as 42, 46, and 41 cells across the three successive biopsies.

Reads were aligned to hg19. Copy-number processing used `CopyKit` version 0.1.4 at revision 9f0d671 [26]. The chromosome-Y-free hg19 VarBin layout comprised 12,167 ordered, approximately 220-kb bins on chromosomes 1–22 and X. All retained cells were jointly segmented with multipcf using *γ* = 40 and segmentation seed 17. Integer states were assigned with scquantum using penalty 25. We then applied nearest-neighbor smoothing with *k* = 4 and repeated the same joint segmentation and integer-state assignment. Biopsy labels were not used in alignment, segmentation, integer calling, smoothing, or cell-quality selection.

The feasibility input was fixed before CTBF reconstruction. It required a mean nearest-neighbor profile correlation of at least 0.90 within the CopyKit quality assessment, in addition to the one-million-read technical threshold. Seventy-nine cells passed both criteria, comprising 28 pretreatment, 16 mid-treatment, and 35 post-treatment cells. This attrition was strongly imbalanced across biopsy times. The retained counts were therefore not used to estimate clone frequencies or the cellular composition of any specimen. Exact integer profiles were collapsed within each biopsy but never between biopsies. This produced 77 biopsy-indexed occurrences and 76 distinct CNP states. One state observed at more than one biopsy consequently retained multiple occurrences.

### 12.2 Absolute-scale ambiguity and analysis boundary

We examined absolute-scale identifiability before computing any CTBF distance or reconstruction. Unsmoothed and *k* = 4 smoothed CopyKit estimates were compared with independent `scAbsolute` estimates at 500-kb resolution for the same 129 technically retained cells [23]. The CopyKit profiles were projected by genomic-overlap weighting onto the scAbsolute autosomal grid. The comparison used 4,680 bins with at least 80% CopyKit coverage. Mean inferred ploidy was 3.052 for scAbsolute, 2.292 for smoothed CopyKit, and 2.240 for unsmoothed CopyKit. The scale disagreement depended on biopsy time. In 107 of 129 cells, the ratio between smoothed CopyKit and scAbsolute ploidy was within 20% of a power of two. The nearest ploidy-derived power-of-two adjustment also minimized profile error among the tested adjustments in 123 of 129 cells. These results leave alternative harmonic scales compatible with the available estimates.

Eleven smoothed CopyKit profiles were constant on the common autosomal grid. All eleven were rejected by the profile-correlation criterion, including eight mid-treatment cells. Including these degenerate profiles would change the retained sample composition. The filtering and scale ambiguities therefore limit interpretation of biopsy-specific cell counts. The available Kim et al. data confirm aneuploid sorting but do not provide a numerical KTN152 DNA index that distinguishes the harmonic solutions [16]. scAbsolute resolves recent whole-genome-duplication and G2 ambiguities through genomic read density when coverage and library design permit it [23]. KTN152 was sequenced as amplified single-end libraries, so this read-density stage was disabled prospectively. The scAbsolute result was therefore retained as a caller sensitivity and not used as ground truth or to rescale CopyKit.

The absolute copy-number scale therefore remained unresolved. For the narrower feasibility application, the 79-cell smoothed CopyKit subset was frozen without reference to CTBF output. The reconstruction input consists of our CopyKit-derived integer CNP estimates. No profile, chromosome, cell, or biopsy was rescaled or selected after reconstruction. Among the 936,859 entries in the 77-occurrence by 12,167-bin matrix, 5,669 (0.61%) were at least 8 and 2,761 (0.29%) exceeded 8. Both thresholds were reached in 70 occurrence profiles and 132 distinct bins, and the maximum estimate was 40. Counts weight each within-biopsy-collapsed occurrence once. The pinned MEDICC2 parser^2^ accepts values through 8 and caps larger values. It therefore cannot preserve the above-8 estimates in its total-copy-number input. These empirical profiles were not submitted to MEDICC2 and were not clipped for CTBF reconstruction. An empirical reference tree is unavailable, so the evaluation concerns computational feasibility and the properties of the inferred representations.

### 12.3 Distance construction and reconstruction

For states *s, t* ∈ *U*, let 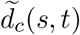 denote the directed `cnp2cnp` cost from *ξ*(*s*) to *ξ*(*t*) restricted to chromosome *c*. The empirical whole-genome dissimilarity was

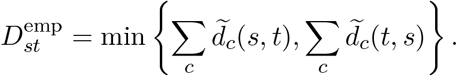

Computing each directed cost separately by chromosome prevents an inferred interval event from crossing a chromosome boundary. The resulting 76 *×* 76 matrix was computed once and supplied unchanged to *M*_*0*_ and *M*_*2*_. Method *M*_*0*_ pooled the 76 distinct states. Method *M*_*2*_ retained the 77 biopsy-indexed occurrences and used the three samples in chronological order.

Both reconstructed graphs satisfied the formal conditions for partial CNP-trees and contained all observed states. Table 16 reports their descriptive dimensions and resource measurements.

The distance stage launched the two directed calculations for each of 23 chromosomes with at most six concurrent workers. Peak memory is the sampled resident memory of the complete process tree. Reconstruction resources were measured in isolated processes on the same 12-core Apple M4 Pro workstation.

The *M*_*0*_ tree had 76 leaves and maximum depth 18. The *M*_*2*_ tree had 66 leaves and maximum depth 14. Its 108 labeled vertices comprised 77 observed occurrences and 31 inferred copy-up occurrences. Relative to the 76 distinct states, these comprise 32 repeated-label occurrences: one additional observed occurrence and 31 inferred occurrences. At the first and second biopsy transitions, it selected radii 21 and 11, respectively. It made 31 copy-up decisions and deferred six exact minimum-distance ties. These quantities describe the operation of *M*_*2*_ on the frozen input. In the empirical figure, each observed-node color is assigned from its biopsy-indexed occurrence rather than its CNP state. The two observations of the recurrent state consequently retain different biopsy colors. Tree edges and vertices are unchanged from the reconstruction.

The inferred tree contains 7 observed ancestor–descendant occurrence pairs between the first and second biopsies, 15 between the second and third, and 24 between the first and third. Each count refers to ordered pairs of observed vertices joined by a nonempty directed path. The illustrated path *v*_*1*_ → *v*_*2*_ → *v*_*3*_ consists of observed occurrences from the first, second, and third biopsies, respectively. Its two edges have dissimilarities 14 and 10, below the corresponding transition radii of 21 and 11. The on-treatment occurrence *v*_*2*_ has eight observed surgical descendants, including *v*_*3*_. The path was chosen for legibility. These relationships are hypotheses conditional on the CNP estimates and reconstruction model, with chronological consistency enforced by the method’s temporal constraint.

### 12.4 NestedBD-Long convergence limitation

We assessed the feasibility of an empirical comparison with NestedBD-Long [17] in a separate exploratory analysis. Following its released input converter, we capped copy numbers at 9 and retained the first bin and every twentieth bin thereafter. The resulting 609-bin profiles were collapsed within, but not between, biopsies, giving 76 biopsy-indexed occurrences (28, 16, and 32) and 75 distinct states. The full-resolution CTBF input above was retained separately. Sampling times for this exploratory comparison were treatment-cycle positions 0, 2, and 6, used as proxies for elapsed clinical time.

We used BEAST 2.6.7 and the NestedBD-Long configuration at revision 3140ac96. Preliminary runs produced nonfinite values for the unused ucldMean parameter. For the runs reported here, we removed its XML state declaration, scale operator, trace-log entry, and the entire joint parameter–tree up–down operator. The biological likelihood, retained prior expression, actual mean clock, and remaining parameter settings were unchanged. The released inference source and binary were unchanged. The XML changes altered the sampled state space and proposal schedule.

Two one-million-iteration technical checks preceded two independently seeded, uninterrupted runs of 80 million iterations, each with four coupled Markov chains. The seeds were 2026092401 and 2026092402. Production samples were recorded every 10,000 iterations. The prespecified burn-in was 10%. Burn-ins of 25% and 50% were examined only as post hoc diagnostic sensitivities. Effective sample sizes (ESS) were computed from each cold-chain trace using the released BEAST LogAnalyser. For between-run tree agreement, a clade was identified by its set of descendant occurrence identifiers. Singleton clades and the complete taxon set were excluded, and zero-length sampled-ancestor branches were retained. A majority clade occurred in more than half of the retained trees in a run.

All numeric trace columns were finite, but low ESS and discordant clade frequencies indicated inadequate mixing (Table 17). The first run underwent a marked posterior shift near 70 million iterations. Discarding the initial half did not resolve the discrepancy. At 50% burn-in, only 21 majority clades were shared between the runs, and the maximum absolute difference in sampled clade frequency was 0.964.

**Table 17:** NestedBD-Long cold-chain diagnostics and between-run clade agreement. Paired entries give run 1 and run 2 in the seed order stated in the text. The 10% burn-in was prespecified.

| Burn-in | Posterior ESS | Tree-height ESS | Majority<br>clades | Shared |
| --- | --- | --- | --- | --- |
| 10% | 8.42 / 31.22 | 11.17 / 235.38 | 55 / 61 | 20 |
| 25% | 6.90 / 35.14 | 8.83 / 174.36 | 56 / 61 | 22 |
| 50% | 4.53 / 644.15 | 6.66 / 136.31 | 55 / 63 | 21 |

The two concurrent production runs took 15.55 and 15.64 hours, with peak resident memory of 239 and 255 MiB, respectively, for the JVM process trees. These measurements cover the production JVM runs, excluding the short trials, controller, and postprocessing. Convergence remained unestablished for this input and configuration under the tested budget. The two runs were retained separately as diagnostic outputs, and their topologies were excluded from the empirical comparison with CTBF.

## Footnotes

1 `cnpsolver.py` at commit `f0b1a19`.

2 MEDICC2 `medicc/io.py`, revision `3f1b8a6`

## Notes

### Competing Interest Statement

The authors have declared no competing interest.

